# Influenza hemagglutinin subtypes have different sequence constraints despite sharing extremely similar structures

**DOI:** 10.64898/2026.01.05.697808

**Authors:** Jenny J. Ahn, Timothy C. Yu, Bernadeta Dadonaite, Caelan E. Radford, Jesse D. Bloom

**Author notes:** These authors contributed equally to this work (Jenny J. Ahn, Timothy C. Yu).

## Abstract

Hemagglutinins (HA) from different influenza A virus subtypes share as little as ∼40% amino acid identity, yet their protein structure and cell entry function are highly conserved. Here we examine the extent that sequence constraints on HA differ across three subtypes. To do this, we first use pseudovirus deep mutational scanning to measure how all amino-acid mutations to an H7 HA affect its cell entry function. We then compare these new measurements to previously described measurements of how all mutations to H3 and H5 HAs affect cell entry function. We find that ∼50% of HA sites display substantially diverged preferences for different amino acids across the HA subtypes. The sites with the most divergent amino-acid preferences tend to be buried and have biochemically distinct wildtype amino acids in the different HA subtypes. We provide an example of how rewiring the interactions among contacting residues has dramatically shifted which amino acids are tolerated at specific sites. Overall, our results show how proteins with the same structure and function can become subject to very different site-specific evolutionary constraints as their sequences diverge.

## Introduction

Proteins often evolve at the sequence level while largely retaining their structures and functions^1,2^. A particularly striking example of this evolutionary dynamic is the hemagglutinin (HA) of influenza virus. There are at least 19 subtypes of HA (numbered H1 to H19), most of which are found in avian influenza viruses with some also found in viruses that have jumped to humans or other mammals^3^, and a handful found only in bat viruses^4^. These HA subtypes have highly conserved structures and share the ability to mediate cell entry via a pH-dependent conformational conversion after binding to receptor^5–7^, which is usually sialic acid although sometimes MHC class II^8,9^. Yet despite this structural and functional conservation, HA subtypes share as little as ∼40% amino-acid sequence identity. This high sequence divergence in the context of a conserved structure and function is driven in part by host immune pressure for antigenic diversification of HA to evade antibodies elicited by prior infections^10,11^. The remarkable extent to which HA maintains its functional properties despite its sequence divergence is illustrated by the fact that two influenza pandemics in the last century were caused by human strains acquiring HAs of different subtypes while retaining most other viral genes: in 1957 a human H1N1 strain acquired a H2 HA with only 67% identity while retaining 5 of the other 7 viral genes, and in 1968 a human H2N2 strain acquired a H3 HA with only 41% identity while retaining 6 of the other 7 genes^12^.

A fundamental question in protein evolution, which is also of public-health relevance in the case of influenza HA, is how this sequence divergence on a nearly fixed structural backbone affects tolerance to further mutations. Amino acids have different propensities for certain secondary structures^13,14^ or hydrophobic versus polar environments^15,16^, and so when the backbone structure is preserved there is expected to be some conservation of the preferences of specific sites for different amino acids. But even on a nearly fixed backbone, sequence divergence can result in epistatic shifts in a site’s amino-acid preferences due to changes in side-chain interactions or subtle shifts in the backbone^17,18^. Experiments have shown that mutations often have similar effects in closely related homologs albeit with occasional epistasis^19–21^, but that the consistency of effects of mutations decays as the homologs become more diverged^22^. Over long evolutionary times, these shifts can lead to coevolution at structurally interacting sites^23–26^.

Here we compare the effects of mutations across subtypes of influenza HA, which as mentioned above exemplifies a protein with high sequence divergence but exceptionally conserved structure and function. To do this, we first use pseudovirus deep mutational scanning^27^ to measure how all mutations to the ectodomain of a H7 HA affect its ability to mediate virion entry into cells. We then compare these measurements to previously generated experimental data for H5 and H3 HAs^28,29^. Overall, we find that the evolutionary divergence of HA across subtypes has led to dramatic changes in mutational tolerance via rewiring of residue interactions. In addition, the new H7 HA deep mutational scanning data reported here provides sequence-function information that can help inform vaccine immunogen design and viral surveillance^28,29^ for an influenza subtype of potential relevance to public health.

## Results

### Phylogenetic and structural comparison of HAs from H3, H5, and H7 subtypes

The HA proteins of influenza A viruses of different subtypes are highly diverged at the sequence level, with amino-acid identities among different subtypes ranging from 36% to 80% (**Fig. 1A,B**). Here we focus on HAs from three subtypes: H3, H5, and H7 (**Fig. 1A**). H3 and H7 HAs have 47% amino-acid identity, while the H5 HA is more diverged from these other two, with identities of 40% and 43% to H3 and H7, respectively (**Fig. 1B**).

**Figure 1.**
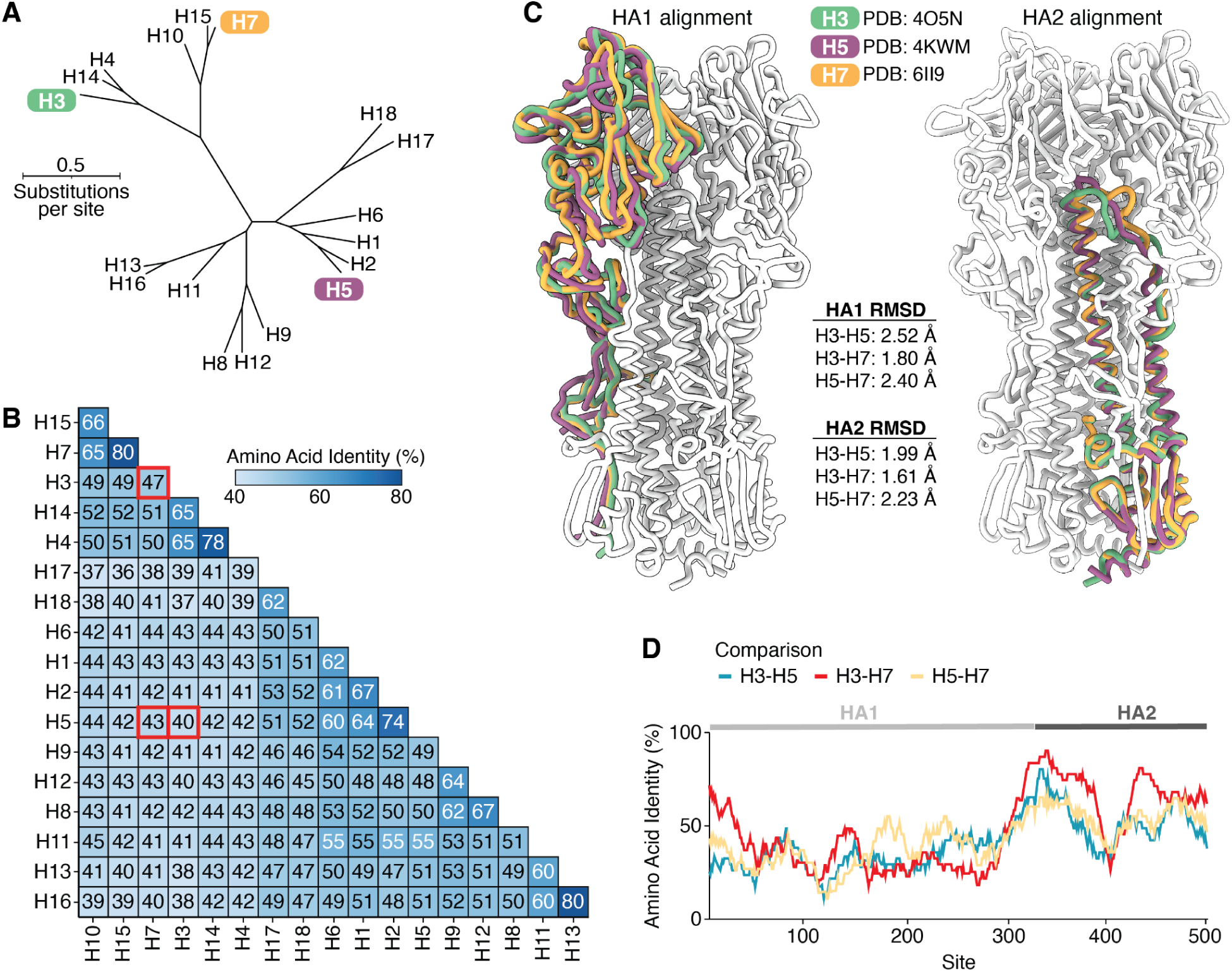
| Phylogenetic and structural comparison of H3, H5, and H7 HAs. **A)** Maximum-likelihood phylogenetic tree of HA protein sequences inferred using IQ-TREE^63^ with the Jones-Taylor-Thornton amino-acid substitution model with rate variation. **B)** Pairwise amino acid identity between the HAs shown in **A**, with red boxes indicating comparisons between the three subtypes that are the focus of the experimental work in this study. **C)** Structural alignments of single HA1 or HA2 monomers of H3 (green), H5 (purple), and H7 (orange) HAs. The rest of the H3 HA trimer (white) is included to provide structural context. HA1 and HA2 were aligned separately due to a shift in the relative orientation of these domains (see **Fig. S1**). The root mean square deviation (RMSD) of carbons between the aligned HA1 and HA2 monomers are also reported. **D)** Pairwise amino acid identities between the different HA subtypes computed along the length of the primary sequence with a sliding window of 30 residues. The HA structures used in this figure are PDB 6II9 (H7)^64^, PDB 4O5N (H3)^65^, and PDB 4KWM (H5)^66^.

Despite their extensive sequence divergence, H3, H5, and H7 HAs have highly conserved structures (**Fig. 1C**). The root mean square deviations of the structurally aligned backbones of the two HA polypeptide chains (HA1 and HA2) are within 2.5 angstroms across experimentally determined structures of HAs from the three subtypes (**Fig. 1C**), although there is some modest shift in the relative orientation of HA1 and HA2 (**Fig. S1A,B**). All three HAs also have conserved biological functions of binding to sialic acid receptors and then mediating membrane fusion at acidic pH^5–7^. Viruses with H3 HAs have circulated in a variety of avian and mammalian species, including humans, dogs, and horses^3^. While avian H3 HAs preferentially bind α2-3 linked sialic acids, some mammalian-adapted H3 strains (including the H3N2 strains that are currently endemic in humans) preferentially bind to α2-6 linked sialic acids^30^. Viruses with H5 and H7 HAs mostly circulate in avian species, but they have caused sporadic infections of humans and other mammalian species^31^, and a H5N1 strain recently became endemic in dairy cattle^32^. Avian H5 and H7 strains preferentially bind α2-3 linked sialic acids, although there are known mutations that can shift this binding preference from α2-3 to α2-6 linked sialic acids^33–37^.

The sequence divergence among H3, H5, and H7 HAs is asymmetrically distributed across the protein (**Fig. 1D**). The fusion peptide spanning sites 330-350 (throughout this paper, we use mature H3 numbering) is the most conserved region, consistent with its key function in mediating membrane fusion. Sites in HA1 tend to be more variable among the HA subtypes than sites in HA2, reflecting HA1’s higher mutational tolerance^38,39^ and positive selection for HA1 mutations that reduce antibody recognition^40,41^.

### Pseudovirus deep mutational scanning measurements of how mutations to an H7 HA affect its cell entry function

The goal of our study is to compare the effects of amino-acid mutations on HA’s biochemical function of mediating virion cell entry across different HA subtypes. We have previously described a pseudovirus deep mutational scanning approach that can be used to measure the effects of all individual amino-acid mutations to HA on its ability to mediate the entry of pseudotyped lentiviral particles into cells^27^ (**Fig. S2A,B**), and applied this approach to H5 and H3 HAs^28,29^. Note that this experimental approach uses lentiviral particles that encode no viral protein other than HA, and are therefore only able to undergo a single round of cell entry—meaning that they are not fully infectious pathogens capable of causing disease, making them a safe tool to study the effects of mutations to HA. Here we extend this previously described approach to measure the effects of mutations to a H7 HA on cell entry in order to create an expanded dataset to facilitate comparison of sequence constraints across HA subtypes.

For these new experiments, we selected the HA of one of the World Health Organization’s candidate H7 vaccine viruses^42^, the Eurasian lineage A/Anhui/1/2013 strain. This strain is from the Eurasian H7N9 lineage that first caused human infections in 2013 in China^43,44^. The human infections were traced to closely related viruses from this same lineage circulating in poultry and live bird markets^45,46^, with poultry infections identified concurrently with the human cases. The H7 HA from the A/Anhui/1/2013 strain is the background for three current candidate vaccine viruses, NIBRG-268, NIIDRG-10.1, and IDCDC-RG33A^47^. The HA from this strain has been shown to bind both the α2-3 linked sialic acids that are the typical receptor for avian-adapted influenza viruses and the α2-6 linked sialic acids that are the typical receptor for human-adapted influenza viruses^48^. The ability of the A/Anhui/1/2013 H7 HA to bind α2-6 linked sialic acids is due primarily to the fact that it has the Q226L mutation^48^, which promotes binding to α2-6 linked sialic acids in multiple HA subtypes^49–51^.

We created duplicate mutant libraries of this H7 HA in the context of pseudoviruses capable of undergoing only a single round of cell entry. The two libraries contained 95% and 96% of all possible amino-acid mutations to the HA ectodomain, respectively. Most of the barcoded HA variants in the libraries contained single amino acid mutations, although a small fraction contained no mutations or multiple mutations (**Fig. S2C-E**). We measured the ability of each barcoded HA variant to mediate pseudovirus entry into cells using previously described approaches^27–29^ (**Fig S2B**), and used global epistasis models^52,53^ to determine the effect of each mutation on cell entry. The effect of each mutation is quantified as the log_2_ of the cell entry of pseudovirus with that mutant HA relative to the unmutated HA, so mutations that do not impact cell entry are assigned effects of zero while mutations that impair entry are assigned negative effects. We measured entry into previously described 293 cells^54,55^ that have been engineered to express only α2-3 linked sialic acids, only α2-6 linked sialic acids, or an equal mix of cells expressing each of the two types of sialic acids. We performed two experimental replicates for each of the two independent pseudovirus mutant libraries, and the measured effects of the mutations on cell entry were highly correlated across both replicates of the same library and between the two independent libraries (**Fig. S3**); throughout this paper we report the median of the measurements across all replicates.

The effects of nearly all mutations to the H7 HA ectodomain on entry into a mix of 293 cells expressing α2-3 linked or α2-6 linked sialic acids are shown in **Fig. 2A** (see also the interactive heatmap at https://dms-vep.org/Flu_H7_Anhui13_DMS/cell_entry.html). As can be seen from this figure, there is wide variation in mutational tolerance across the HA protein, with some sites being quite tolerant of mutations whereas at other sites nearly any mutation strongly impairs HA’s cell-entry function. Functional constraint was generally higher in HA2 than HA1, especially in regions of HA2 like the fusion loop (sites 330-350) that are critical for mediating HA’s cell entry function (**Fig. 2A**). However, many regions of HA1, especially near the top of the globular head of the protein, are quite tolerant of mutations (**Fig. 2A,B**). Overall, these results are in line with prior deep mutational scanning studies of HA from other influenza A subtypes, which have found the most constraint in regions of HA2 involved in membrane fusion, and higher mutational tolerance in many portions of the HA1 globular head including the epitopes most commonly targeted by neutralizing antibodies^28,38,56–58^.

**Figure 2.**
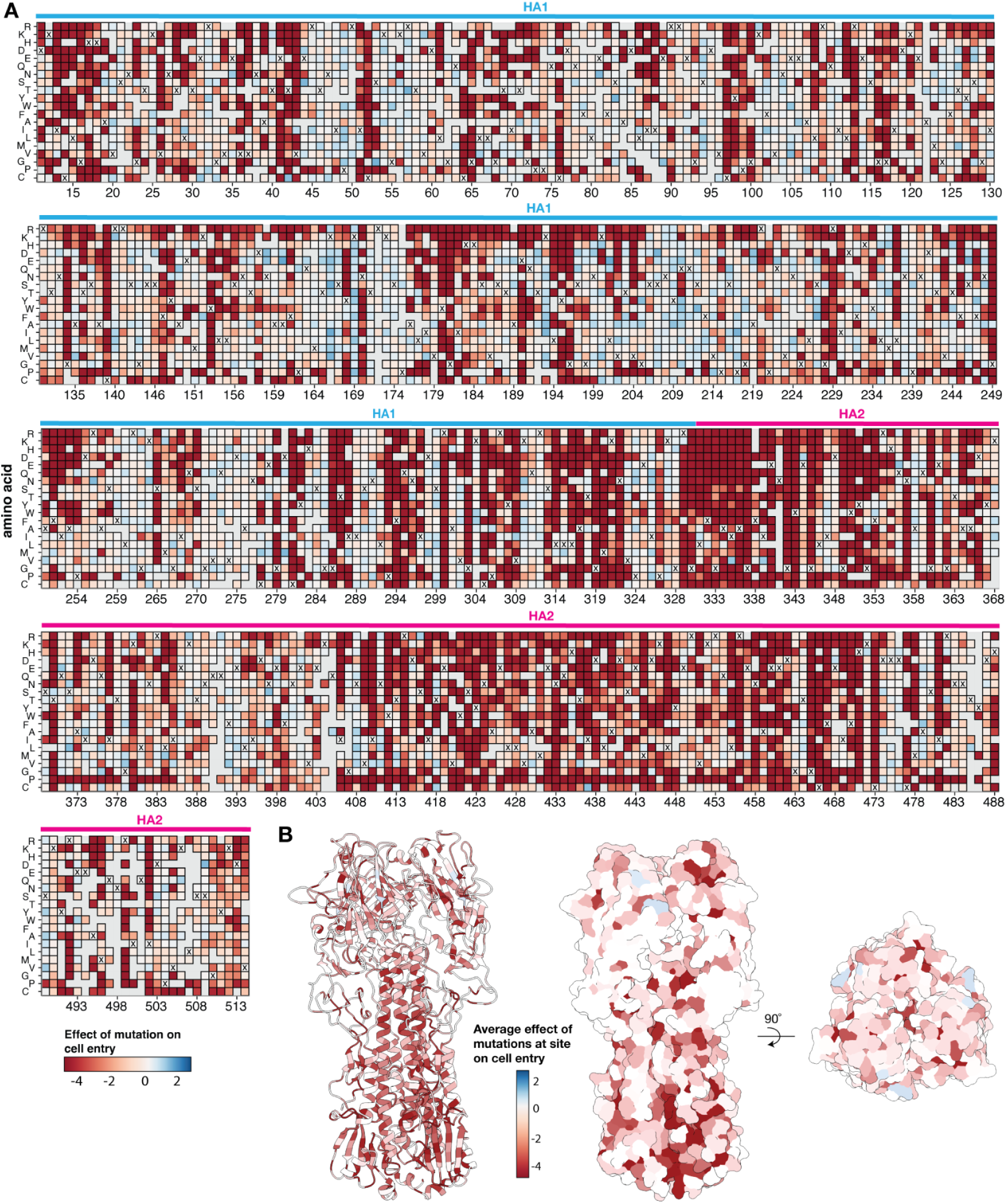
| Effects of mutations to the ectodomain of H7 HA on its ability to mediate entry into a mix of cells expressing α2-3 and α2-6 linked sialic acids. **A)** Heatmap showing the effects of mutations on HA-mediated cell entry. Negative cell entry effects (red) indicate impaired cell entry relative to the unmutated HA. Gray indicates mutations that were not reliably measured in our experiments, and for each site the X indicates the amino-acid identity in the unmutated A/Anhui/1/2013 HA. See https://dms-vep.org/Flu_H7_Anhui13_DMS/cell_entry.html for an interactive version of this heatmap that also shows effects of mutations on entry into 293 cells expressing only α2-3 or only α2-6 linked sialic acids. In this figure and throughout the rest of this paper we use H3 sequence numbering^67^. **B)** The structure of the A/Anhui/1/2013 H7 HA^64^ (PDB 6ii9) colored by the mean effects of all mutations at that site on cell entry.

The effects of H7 HA mutations on entry into 293 cells expressing only α2-3 linked or only α2-6 linked sialic acids were very similar to the effects of mutations on entry into the mix of cells expressing α2-3 linked and α2-6 linked sialic acids (**Fig. S4A**). There were only a few sites where the effects of mutations were appreciably different in the α2-3 linked only or α2-6 linked only sialic acid cells; these were mostly sites in the receptor-binding loops known to be important for receptor binding specificity such as 193, 220, and 226 (**Fig. S4A**). The reason that the effects of mutations are so similar on entry in both types of cells is likely because most HA mutations affect entry by modulating HA folding and membrane fusion rather than receptor-binding, and because the A/Anhui/1/2013 H7 HA used in our experiments is already capable of binding both α2-3 linked and α2-6 linked sialic acids due to the fact that it contains the Q226L mutation^48^. To confirm this latter point, we created pseudovirus with the L226Q reversion and measured its entry into the 293 cells expressing only α2-3 linked or only α2-6 linked sialic acids (**Fig S4B**). Pseudovirus with the unmutated A/Anhui/1/2013 H7 HA entered both cells similarly, but pseudovirus with the Q226L revertant of this protein were better at entering the α2-3 linked sialic acid expressing cells. As additional controls, we confirmed pseudovirus expressing an avian H5N1 HA was better at entering the α2-3 linked sialic acid expressing cells, while pseudovirus expressing a human H3N2 HA was better at entering the α2-6 linked sialic acid expressing cells (**Fig S4B**). Overall, these results show that the effects of most mutations to the H7 HA on its cell entry function are the same regardless of the type of sialic acid expressed on the target cells, although a small number of mutations in the receptor binding loops affect entry in a way that depends on the sialic acid expressed on the target cells.

### Comparing amino-acid preferences across H3, H5, and H7 HAs

We next compared the H7 HA deep mutational scanning dataset described above to previously published H3 and H5 datasets^28,29^ to assess how the effects of amino-acid mutations on HA’s cell entry function differ across these subtypes. To make these comparisons, we converted the measured effects of mutations on cell entry to normalized values quantifying the preference of each site for each amino acid. Quantitatively, the preference π_𝑟,𝑎_ of site *r* for amino acid *a* is defined as proportional to the exponential of the effect 𝑥_𝑟,𝑎_ of that mutation on cell entry, namely as 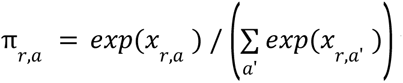 where the sum over *a’* is taken over all amino-acid identities. If all amino-acid mutations except the wildtype identity are highly deleterious at a site then only that wildtype amino acid is preferred; when many mutations are well tolerated at a site then many amino acids have similar preferences (**Fig. 3A**). There are two advantages of converting the measured effects of mutations to amino-acid preferences for comparing across subtypes. First, unlike the mutation effects, the amino-acid preferences at a site are not defined relative to its wildtype amino-acid identity and so facilitate direct comparison across H3, H5, and H7, which often have different amino acids at the same site (**Fig. 1B**). Second, quantitative comparisons using amino-acid preferences emphasize differences in which amino acids at a site are well tolerated, whereas quantitative comparisons using directly measured mutation effects also emphasize less relevant differences between moderately and strongly deleterious amino acid identities at a site. Given the experimentally measured set of amino-acid preferences at a site in two different HA subtypes, we quantified the difference in evolutionary constraint at the site by the Jensen-Shannon divergence. This divergence is high when mutations at a site have dramatically different effects between HA subtypes, and low when mutations have similar effects in both HA subtypes (**Fig. 3B**).

**Figure 3.**
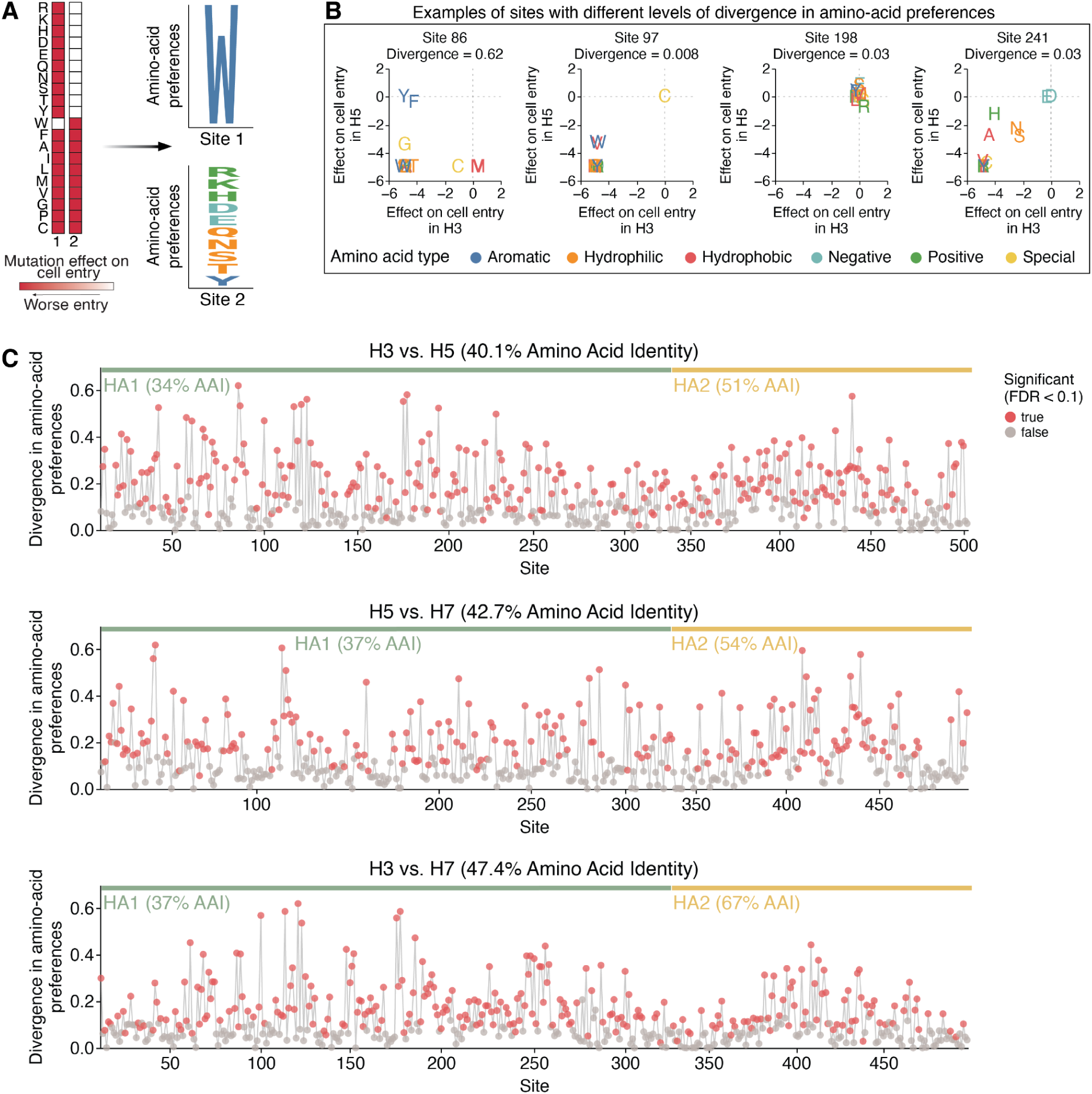
| Sequence constraints differ at many sites among H3, H5, and H7 Has. **A)** Mutation effects on cell entry at a site are converted into amino-acid preferences, and the amino-acid preferences, which can be plotted as logo plots where the height of the letter is proportional to the preference of that site for that amino acid. **B)** Examples of sites with different Jensen-Shannon divergence in amino-acid preferences between HA subtypes. The scatter plot shows the effect of mutating to each amino acid in each of the two subtypes, with the wildtype identity assigned an effect of zero. **C)** Divergence in amino-acid preferences at each site in HA between H3-H5, H5-H7, and H3-H7. Red points indicate sites where preferences have significantly diverged (false discovery rate < 0.1) between HA subtypes. See https://jbloomlab.github.io/ha-preference-shifts/htmls/combined_interactive_lineplots.html for an interactive version of this plot.

Many HA sites display high divergence in amino-acid preferences among subtypes (**Fig. 3C**). Overall, 58% of sites exhibited significant divergence in amino-acid preferences between H3-H5, 49% of sites between H5-H7, and 54% of sites between H3-H7 (FDR < 0.1, see red points in **Fig. 3C**). At those significantly diverged sites, HAs of different subtypes often preferred distinct amino acid types (**Fig. 3B**, left panel), demonstrating how the biochemical determinants of mutational tolerance differs across HA subtypes. Although the experimental measurements of the effects of mutations on cell entry for the different HA subtypes were performed in different target cells (a mix of 293 cells expressing α2-3- and α2-6-linked sialic acids for H7, MDCK-SIAT1 for H3^29^, and 293T for H5^28^) that express varying ratios of α2-3- and α2-6-linked sialic acids, the use of different target cells is not a major contributor to the divergence in amino-acid preferences across subtypes. This fact is demonstrated by observing that there is very little divergence between the amino-acid preferences of the H7 HA between measurements made using 293-α2-3 versus 293-α2-6 cells (**Fig. S5A,B**), with only a few sites in the receptor-binding pocket exhibiting modest differences in amino-acid preferences between cell types. This result indicates that most of the constraints on mutations are due to their effects on protein folding, stability, and fusion function—which are conserved across cell types. Only at a small handful of sites directly involved in receptor binding are there cell-type specific differences in amino-acid preferences. Collectively, these results suggest the evolutionary constraints on cell entry are dramatically different across H3, H5, and H7 HAs due to large changes in sequence that alter the preferences for specific amino acids at different sites due to their impacts on fundamental aspects of HA protein folding and function.

### Divergence in amino-acid preferences across structural and functional regions of HA

We next examined if divergence in amino-acid preferences differs across the two polypeptide chains of HA: HA1 and HA2. HA1 contains the antigenic regions that are under positive selection from antibodies, while HA2 comprises the stem region that is under purifying selection to maintain HA stability. For the H3-H7 comparison, sites in the HA2 domain showed lower divergence than those in the HA1 domain (*p* < 0.001, two-sided Mann-Whitney-U test, **Fig. S6A**). In contrast, no significant differences in divergence were observed between HA1 and HA2 for the H3-H5 and H5-H7 comparisons. This pattern reflects the higher amino acid conservation (67%) of HA2 between H3-H7 versus H3-H5 and H5-H7 (**Fig. 3C**), which was likely maintained due to strong purifying selection against mutations at sites in the HA stem. In general, sites where amino acids are conserved between HA subtypes are typically mutationally intolerant in both subtypes (**Fig. S6B**). However, the finding that there is no significant difference in divergence across HA1 and HA2 between H3-H5 and H5-H7 indicates that while the networks of interacting residues in HA2 are more preserved than those for HA1 between H3-H7, they have diverged similarly in HA1 and HA2 over the broader evolutionary split between group 1 and group 2 HAs.

Finally, to explore how divergence differs across functional domains in HA, we compared divergence across the receptor-binding pocket and antigenic regions (**Fig. S6C**). In contrast with the comparison across HA1 and HA2 regions, phylogenetic relatedness among HAs does not always underlie divergence at these functional domains. In the 190-helix of the receptor-binding pocket, which overlaps with antigenic epitope B (H3 classification), sites showed the highest median divergence in amino-acid preferences in the H3–H7 comparison relative to the H3–H5 and H5–H7 comparisons (**Fig. S6C**). In this case, divergence may instead reflect distinct selective pressures in the human and avian reservoirs that drove adaptations in receptor binding and antigenic escape. We suggest that a common pressure to maintain receptor specificity in birds convergently shaped the mutational tolerance of the 190-helix and antigenic epitope B for H5 and H7, whereas pressure for changes in receptor specificity and antibody escape in humans drove the divergence in these regions for H3.

### Factors associated with divergence in amino-acid preferences among HA subtypes

We sought to determine what specific factors were associated with high divergence in the amino-acid preferences of the same site across HA subtypes. Sites where the wildtype amino acid identity changed between HAs tended to show significantly higher divergence compared to sites where the wildtype amino acids are conserved (*p* = 0.07 for H3-H5; *p* < 0.001 for H3-H7; *p* < 0.001 for H5-H7; two-sided Mann-Whitney-U test, **Fig. 4A**). This difference likely reflects the fact that conserved sites usually cannot tolerate most mutations across HAs (**Fig. S6B**). However, even sites with conserved wildtype amino-acid identities sometimes have different preferences between HA subtypes (**Fig. 4A**). We therefore examined whether additional structural or biochemical features beyond conservation could help predict a site’s divergence.

**Figure 4.**
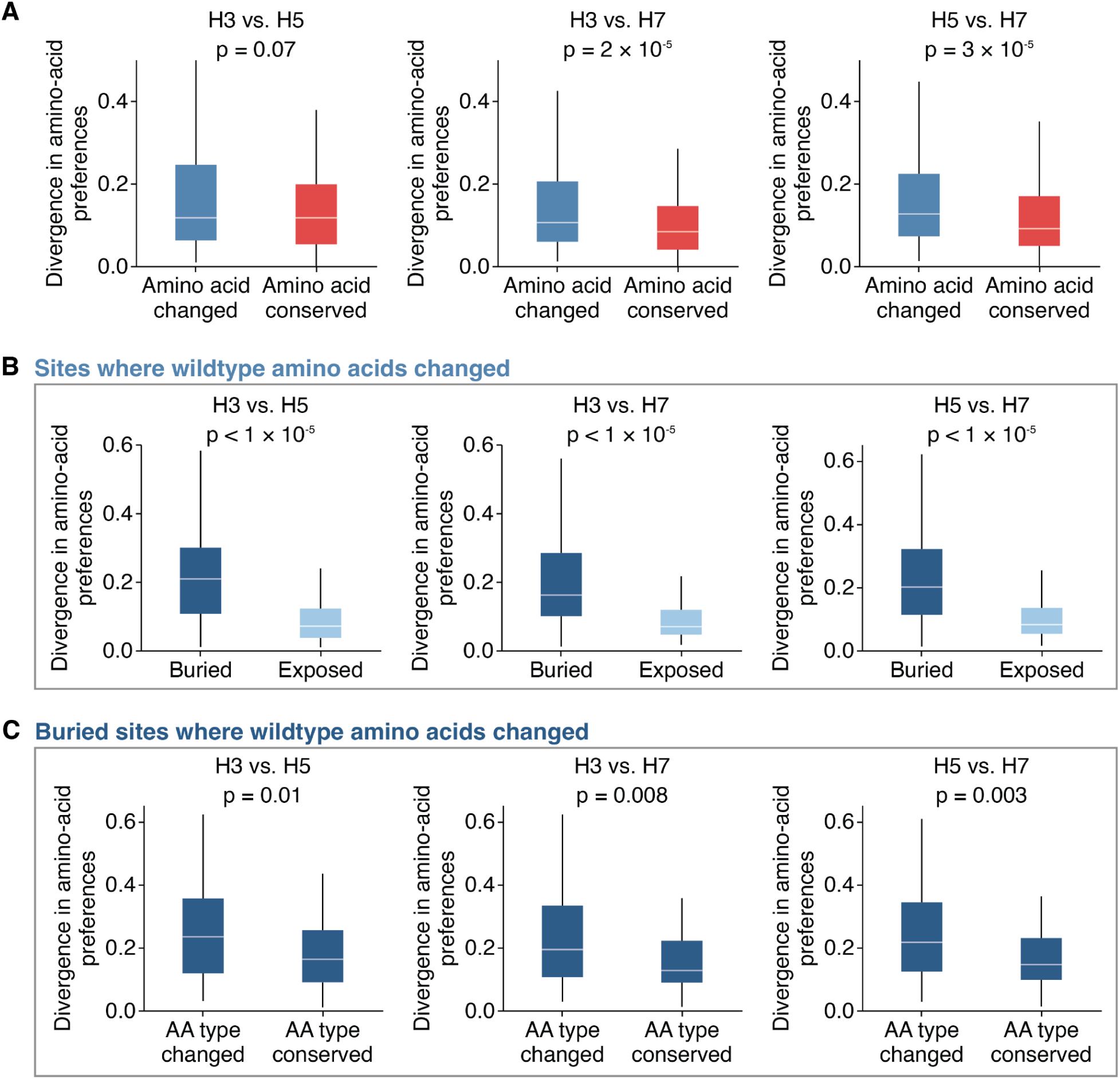
| Sites with divergent amino-acid preferences across HAs tend to be buried and have different wildtype amino-acid types. **A)** Divergence in amino-acid preferences at sites where the wildtype amino-acid identity is conserved or changed between HA subtypes. **B)** Among sites where the wildtype identity changed, buried sites show significantly higher divergence in amino-acid preferences compared to exposed sites. Buried sites are defined as having a relative solvent accessibility < 0.2. **C)** Among buried sites where the wildtype amino-acid identity changed, sites where the wildtype amino-acid type is different (e.g., hydrophobic versus hydrophilic) show significantly higher divergence compared to sites where the amino acid type is conserved. Two-sided Mann-Whitney-U test was used for significance testing.

There is no correlation between the distance of Cα backbone atoms in structurally aligned HAs and divergence (**Fig. S7**), indicating structural shifts in the backbone residues between HA subtypes is not a major driver of divergence in amino-acid preferences. However, among sites where the wildtype amino-acid identity changed between HAs, sites that are buried (relative solvent accessibility ≤ 0.2) showed significantly higher divergence in amino-acid preferences compared to sites that are surface exposed (*p* < 0.001 for all comparisons, two-sided Mann-Whitney-U test, **Fig. 4B**). In contrast, among sites where the wildtype identity is conserved between HAs, solvent accessibility does not significantly explain divergence (*p* > 0.1 for all comparisons, two-sided Mann-Whitney-U test, **Fig. S8B**). These patterns arise because conserved sites tend to be more mutationally intolerant regardless of solvent accessibility (**Fig. S6B**), whereas variable sites that are exposed tend to be more mutationally tolerant in both HA backgrounds compared to variable sites that are buried (**Fig. S8C**). For example, the wildtype amino acids at the exposed site 173 (relative solvent accessibility = 0.85) are Q, R, and K in H3, H5, and H7, respectively, and all measured mutations are tolerated in all backgrounds, resulting in low divergence in amino-acid preferences (**Fig. S8D**). Together, these data suggest that among variable sites where the wildtype amino acid changed, the ones with highly diverged amino-acid preferences are typically buried within the protein.

Yet while the most divergence in amino-acid preferences tends to occur at buried sites with different wildtype amino acids between subtypes, divergence still ranges widely at such sites (**Fig. 4B**). What accounts for this variation? Again, structural deviation of the protein backbone at these sites is not strongly associated with divergence amino-acid preferences (*r* < 0.1 for all comparisons, **Fig. S7**). However, buried sites where the wildtype amino-acid identity differs in biochemical type across subtypes show significantly higher divergence compared to sites where the wildtype amino acids are of the same type (*p* ≤ 0.01 across all comparisons, two-sided Mann-Whitney-U test, **Fig. 4C**). Therefore, the type of amino acid at a buried site where the wildtype identity changed across subtypes is indicative of divergence in amino-acid preferences.

The high divergence of amino-acid preferences at buried sites with different wildtype identities across HA subtypes can arise from fundamental rewiring of how sets of residues interact, even when the backbone structure has not changed. For example, the wildtype amino acids at the buried sites 123/176/178 are E/K/Y in H3, I/L/I in H5, and M/A/I in H7. The amino-acid preferences at these three sites are fairly similar between H5 and H7 HAs, but both are sharply diverged from H3 HA (**Fig. 5A**, **Fig. S9**). This high divergence of amino-acid preferences is likely because sites 123/176/178 form a hydrogen bond network in H3 HAs, but this network is absent in H5 and H7 HAs, which instead have a more hydrophobic environment in this region (**Fig. 5B**). Collectively, the amino-acid preferences and protein structure show how evolution has rewired the constraints in this region: H3 relies on a highly constrained hydrogen bond network to maintain cell entry function, while H5 and H7 rely on a hydrophobic environment that does not require hydrogen bonds at all.

**Figure 5.**
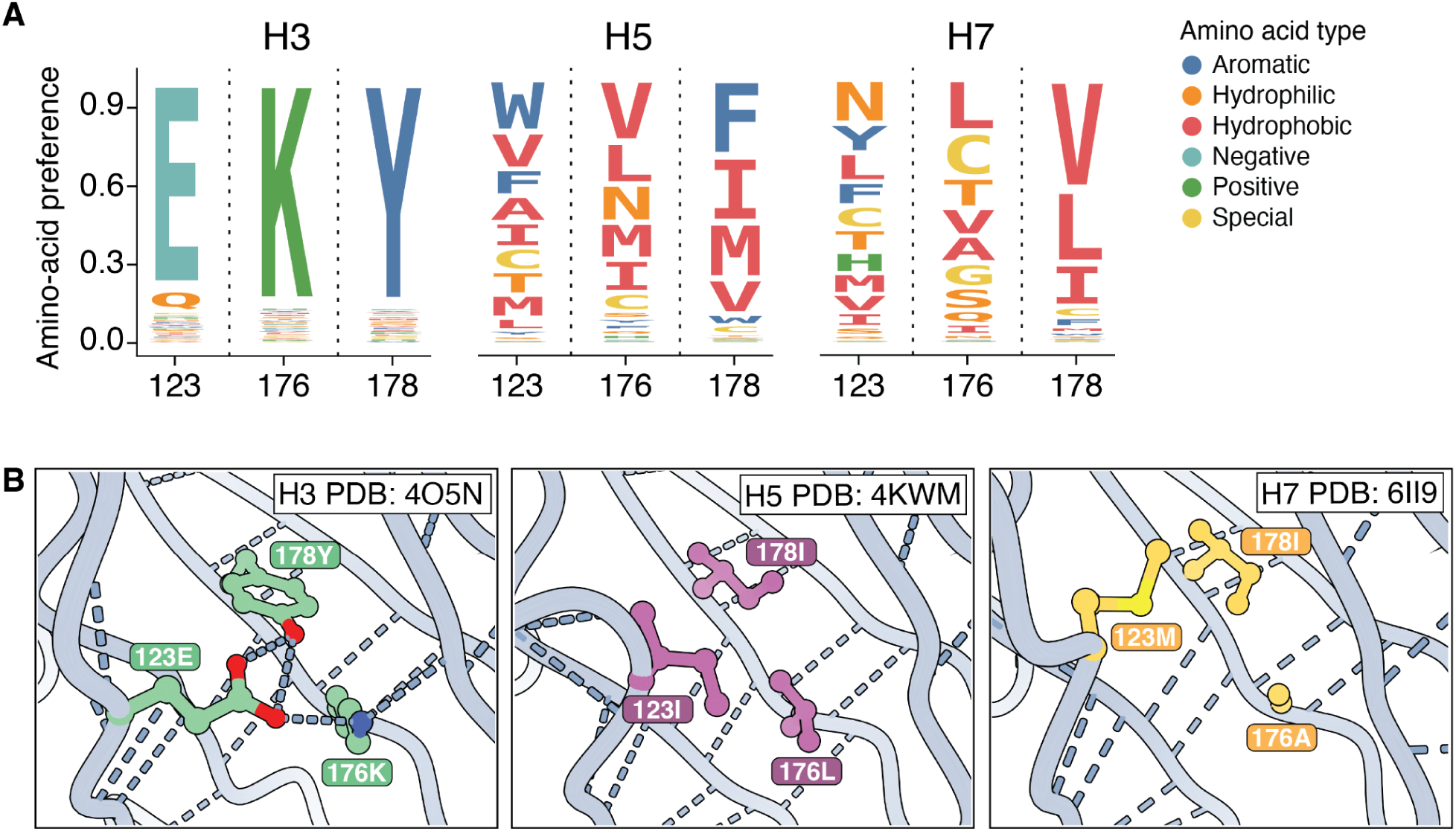
| Example of contacting sites where interactions have been rewired from a hydrogen bond network in H3 HA to a hydrophobic environment in H5 and H7 HAs. **A)** Logoplots showing preferences for amino acids at buried sites 123, 176, and 178 in H3, H5, and H7. The height of each letter is proportional to the amino-acid preference measured by deep mutational scanning. Each amino acid is colored by its biochemical type. The amino-acid preferences at these sites are relatively similar between H5 and H7, but both are highly diverged from H3 HA. **B)** Sites 123, 176, and 178 form a hydrogen bond network in H3, but are part of a hydrophobic environment in H5 and H7 HAs.

## Discussion

Here we have measured how nearly all amino-acid mutations to the ectodomain of a H7 HA affect its cell entry function, and compared the results to previously generated data for H3 and H5 HAs. We find that the effects of mutations at many sites have substantially diverged among the HA subtypes despite the proteins having virtually superimposable backbone structures and mediating the same biochemical function. This divergence in site-specific amino-acid preferences is distributed across HA’s sequence and structure, with the greatest divergence being at buried sites where the wildtype amino-acid type differs between subtypes. In some cases, the highly diverged amino-acid preferences are due to clear rewiring of how the neighboring residues interact even though the protein backbone structure remains unchanged.

It is interesting to contrast the highly diverged amino-acid preferences of different HA subtypes reported here to recent studies that have extensively examined the effects of mutations to the SARS-CoV-2 spike for different viral variants^20,21,52,59–61^. The spikes of even the most diverged SARS-CoV-2 variants are far more similar at the sequence level than the different HA subtypes studied here (current SARS-CoV-2 variants diverged over ∼6 years, whereas HA subtypes diverged over many millenia), and most sites in SARS-CoV-2 spike have similar amino-acid preferences albeit with some examples of epistatic shifts in mutational effects^20,21,52,59–61^. Furthermore, most of the strong epistastic shifts in amino-acid preferences that have occurred during the short-term evolution of the SARS-CoV-2 spike involve sites that affect receptor binding^20,21,59–61^, whereas the most striking changes in amino-acid preferences across the highly diverged HA subtypes involve residues buried in the protein structure that are more involved in folding and stability than receptor binding. This contrast is consistent with recent work on other proteins suggesting that over long evolutionary time frames, pervasive shifts in site-specific amino-acid preferences across a protein accumulate due to many small epistatic interactions among fixed mutations^22^.

It is important to note that our experiments only measure how mutations affect HA-mediated cell entry, which may not fully capture all the ways that HA contributes to viral fitness. In particular, pseudotyped lentiviral particles are not identical in morphology to actual influenza virions, and HA can contribute to transmissibility and immune recognition in ways beyond its ability to mediate cell entry. Therefore, our findings should be interpreted specifically in terms of how mutations affect HA’s cell entry function, which is related to but not identical to its contributions to viral fitness.

Our results have implications for how large-scale sequence-function measurements can inform viral surveillance and vaccine immunogen design. One of the rationales for characterizing the effects of mutations to viral proteins is to provide information that can help interpret the consequences of changes that are observed during genomic surveillance of ongoing viral evolution. Indeed, this rationale is why we focused the new experimental work in this study on H7 HA, since that subtype is thought to pose a potential risk to humans^62^. But of course viruses themselves are always evolving, and so the protein sequence that is the subject of any particular experimental study is unlikely to be identical to that of future viruses of public-health interest. Our results underscore the limits to the extent that experimental measurements made in one genetic background can predict the impacts of mutations in another genetic background. Therefore, reliable prediction of mutation effects across an entire family of a highly diverged but structurally and functionally conserved protein such as HA cannot be achieved simply by large-scale experimental studies—although such studies can be highly predictive for closely related variants of a protein as might be found in the viral strains circulating in humans at any given time.

## Acknowledgements

We thank Hugh Haddox for helpful comments on the manuscript. This work was supported in part by the NIH/NIAID under R01AI165821 and 75N93021C00015. TCY was supported by a NSF Graduate Research Fellowship (DGE-2140004). This research was also supported by the Genomics & Bioinformatics Shared Resource (RRID: SCR_022606), the Flow Cytometry Shared Resource (RRID: SCR_022613) of the Fred Hutch/University of Washington/Seattle Children’s Cancer Consortium (P30 CA015704) and by Fred Hutch Scientific Computing, NIH grants S10-OD-020069 and S10-OD-028685. JDB is an Investigator of the Howard Hughes Medical Institute. This paper is the result of funding in whole or in part by the National Institutes of Health (NIH). It is subject to the NIH Public Access Policy. Through acceptance of this federal funding, NIH has been given a right to make this paper publicly available in PubMed Central upon the Official Date of Publication, as defined by NIH.

## Competing interests

JDB consults for Apriori Bio, Invivyd, Pfizer, GSK, and the Vaccine Company. JDB and BD are inventors on Fred Hutch licensed patents related to the deep mutational scanning of viral proteins.

## Supplemental figures

**Figure S1.**
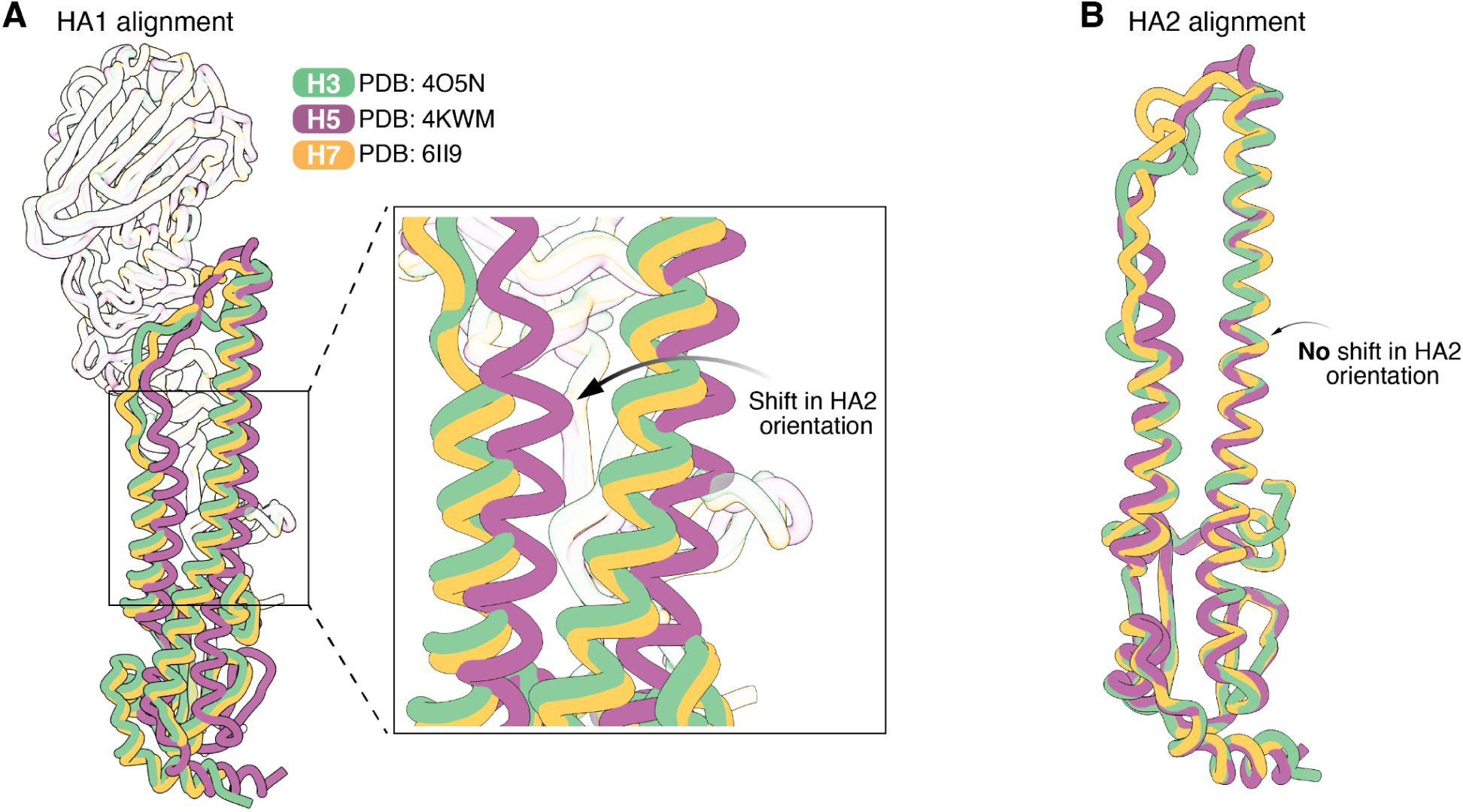
| The relative orientations of HA1 and HA2 are shifted in H5 HA relative to H3 and H7 HAs. **A)** Structural alignment of the HA1 domain (same as left structure in Fig. 1C) with the HA2 domains from H3 (green), H5 (purple), and H7 (orange) HAs. Due to a tilt in the orientation of the HA1 and HA2 domains, the HA2 domain of H5 becomes shifted relative to where the HA2 domains of H3 and H7 are located. **B)** Aligning the HA2 domain alone (same as right structure in Fig. 1C) reveals the folds in HA2 are actually highly conserved across H3, H5, and H7.

**Figure S2.**
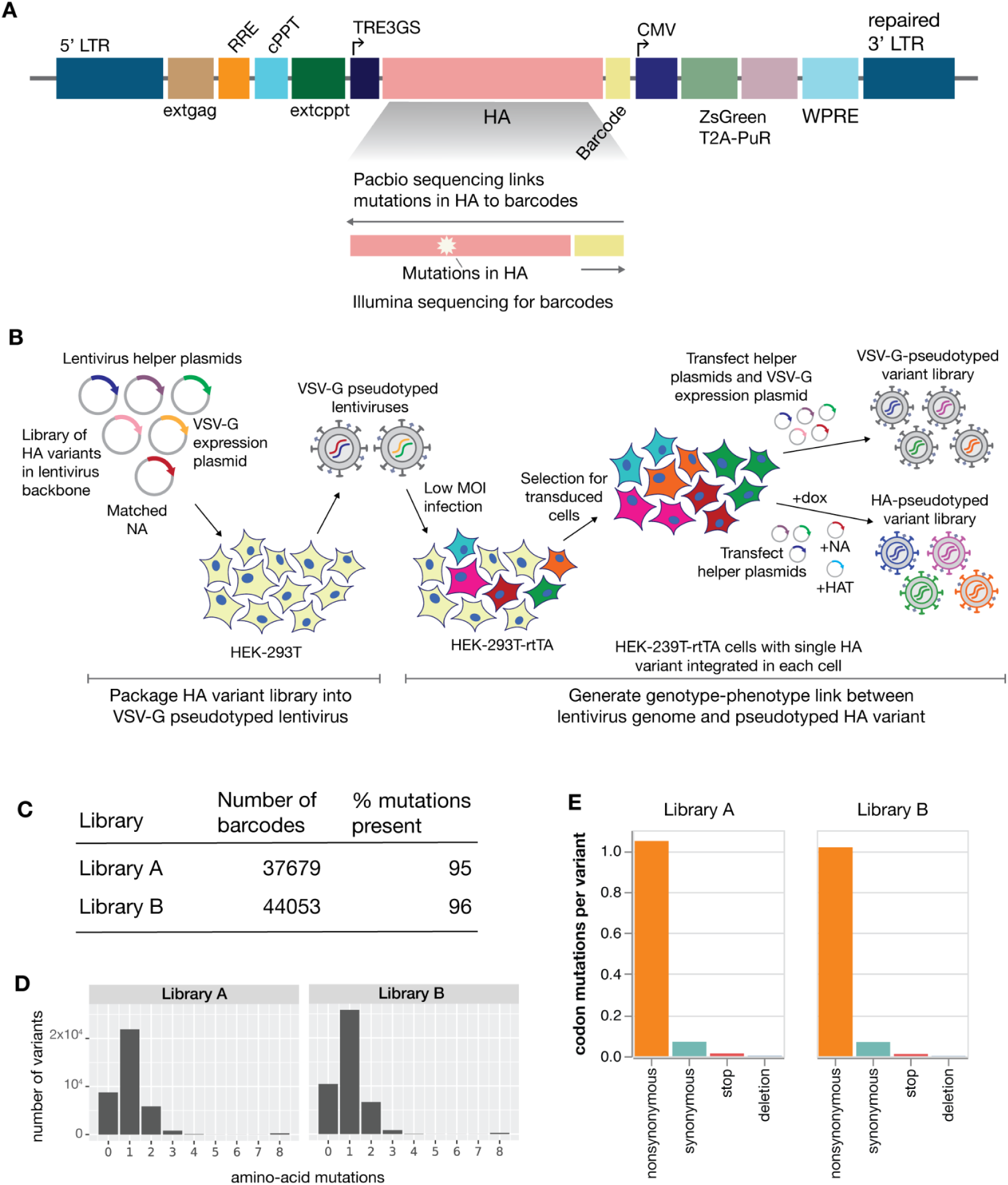
| Overview of pseudovirus deep mutational scanning of H7 HA. **A)** The pseudovirus deep mutational scanning uses a lentiviral backbone encoding the HA protein followed by an identifying 16-nucleotide barcode under the control of an inducible promoter. The lentiviral backbone has a full 3’-LTR so that integrated proviruses can be reactivated by transfection of appropriate helper plasmids. **B)** Schematic of pseudovirus deep mutational scanning experiments. We create a library of mutants of the HA in the context of the barcoded-HA expressing lentiviral backbone plasmid. Cells are transfected with this HA-encoding lentiviral backbone along with helper plasmids encoding the other proteins needed to produce lentiviral particles (Gag-Pol, Tat, Rev), a plasmid encoding the matched N9 NA, and a plasmid encoding VSV-G. The resulting lentiviral particles, which lack a genotype-phenotype link and express VSV-G on their surface, are the infected into rtTA expressing 293T cells at low multiplicity of infection, and transduced cells are infected so that each cell encode a unique HA mutant as a provirus in its genome. These cells are then transfected with the helper plasmids and NA-encoding plasmids to produce pseudoviruses that express unique HA mutants. Cells are also transfected with VSV-G to produce control pseudoviruses that are not dependent on HA to infect cells. To measure cell entry of each HA variant, we use sequencing to compare the efficiency of that barcoded lentiviral particle at entering cells in the presence or absence of VSV-G (particles always enter cells when VSV-G is expressed, but otherwise require a functional HA for cell entry). Note that these lentiviral particles encode no proteins other than HA, and so are not capable of undergoing multicycle replication and so provide a safe way to study the effects of mutations to HA. **C)** Number of uniquely barcoded HA variants and the fraction of all possible HA ectodomain amino-acid mutations that are present in at least one barcoded variant in each of the two independent H7 HA pseudovirus libraries. **D)** Distribution of amino-acid mutations per barcoded HA variant in each library. **E)** Average number of nonsynonymous, synonymous, stop-codon, and deletion mutations per barcoded HA variant.

**Figure S3.**
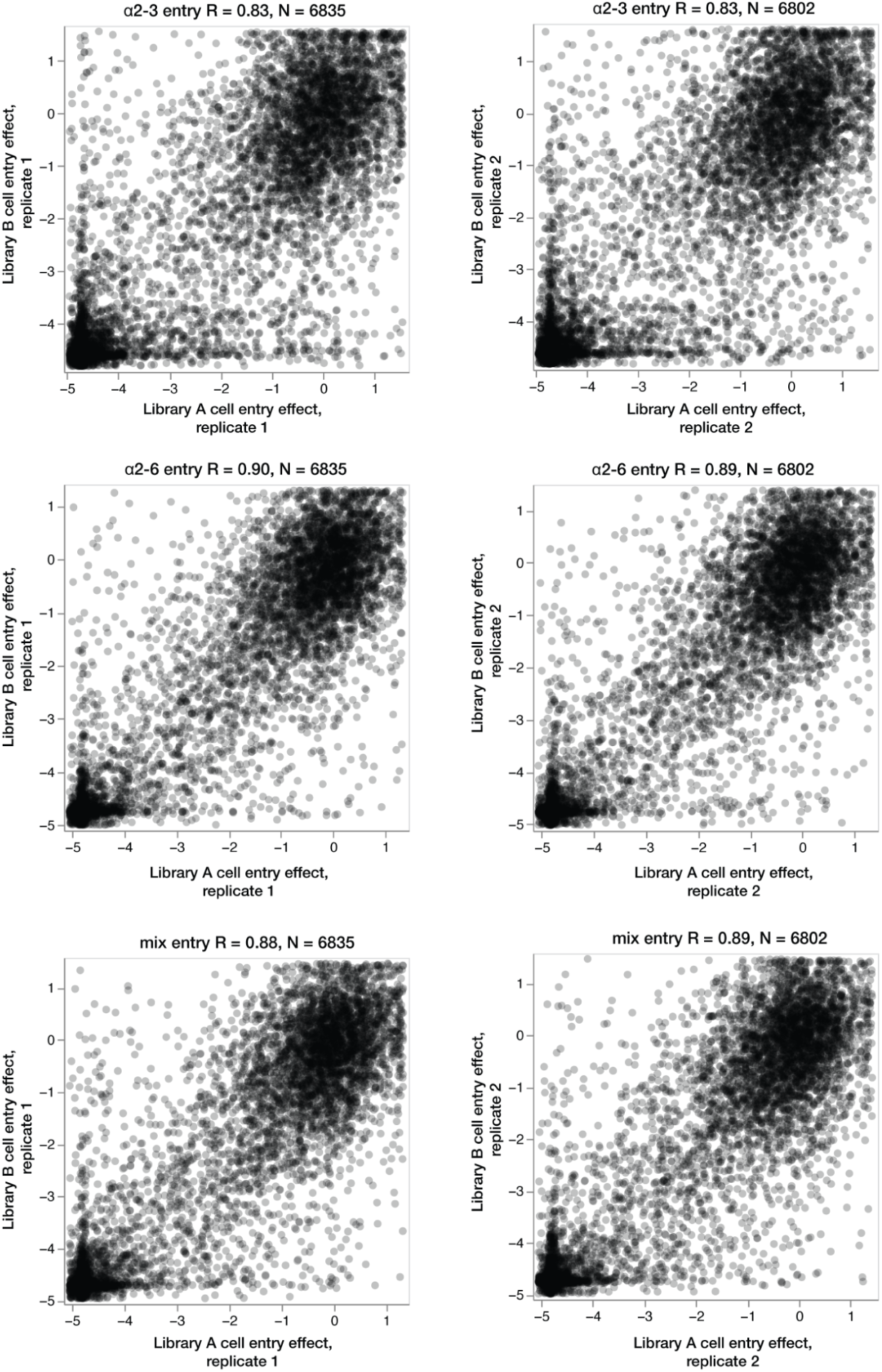
| H7 HA cell entry effects measured using independent deep mutational scanning libraries are highly correlated. Correlation of the effects of H7 HA mutations on cell entry as measured with independent pseudovirus libraries on 293 cells expressing α2-3 linked sialic acids (top row), α2-6 linked sialic acids (middle row), or a mix of both cells (bottom row). Each point represents the effect of a mutation measured in each replicate. For each of the two libraries, two technical replicates were performed. Each panel shows the correlations between two independent libraries (Library A and B) for each technical replicate (replicate 1 or 2). R is the Pearson correlation and N is the number of mutations measured in both pairs of measurements.

**Figure S4.**
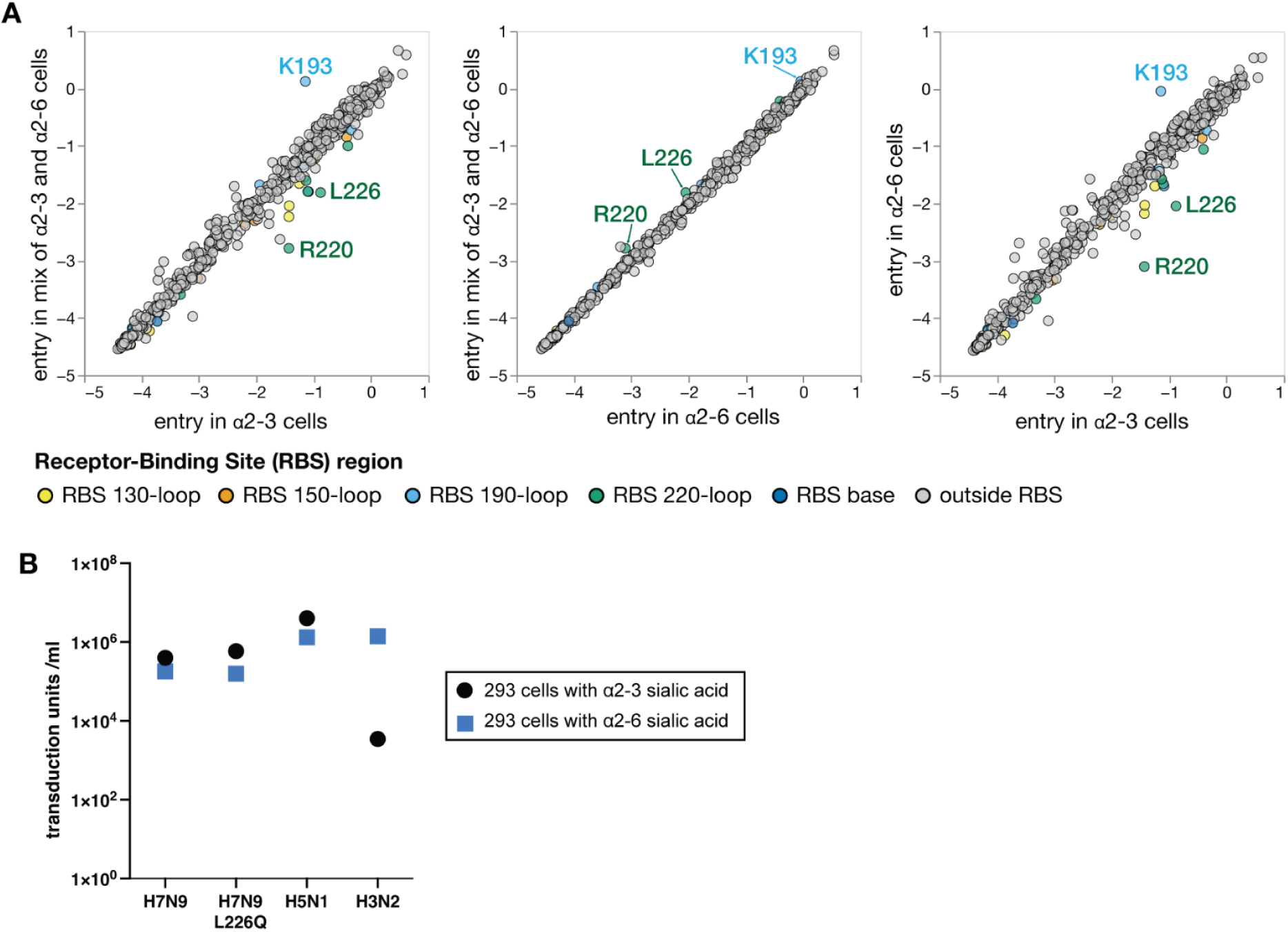
| Effects of mutations to the H7 HA on entry in 293 cells expressing only α2-6 or only α2-3 linked sialic acids. **A)** Scatter plots showing the average effects of mutations at each site on entry into 293 cells expressing only α2-3 linked sialic acids, only α2-6 linked sialic acids, or an equal mix of the two cells. Sites are colored by whether they are part of the receptor-binding site (RBS). See https://dms-vep.org/Flu_H7_Anhui13_DMS/cell_entry.html for interactive versions of these plots. **B)** Titers on 293 cells expressing only α2-3 linked sialic acids or only α2-6 linked sialic acids of lentiviral particles pseudotyped with the HA and NA of A/Anhui/1/2013 (H7N9) strain, the same H7N9 HA and NA but with the L226Q mutation in HA, the HA and NA of the avian A/American Wigeon/South Carolina/USDA-000345-001/2021 (H5N1) strain, or the HA and NA of the human A/Massachusetts/18/2022 (H3N2) strain.

**Figure S5.**
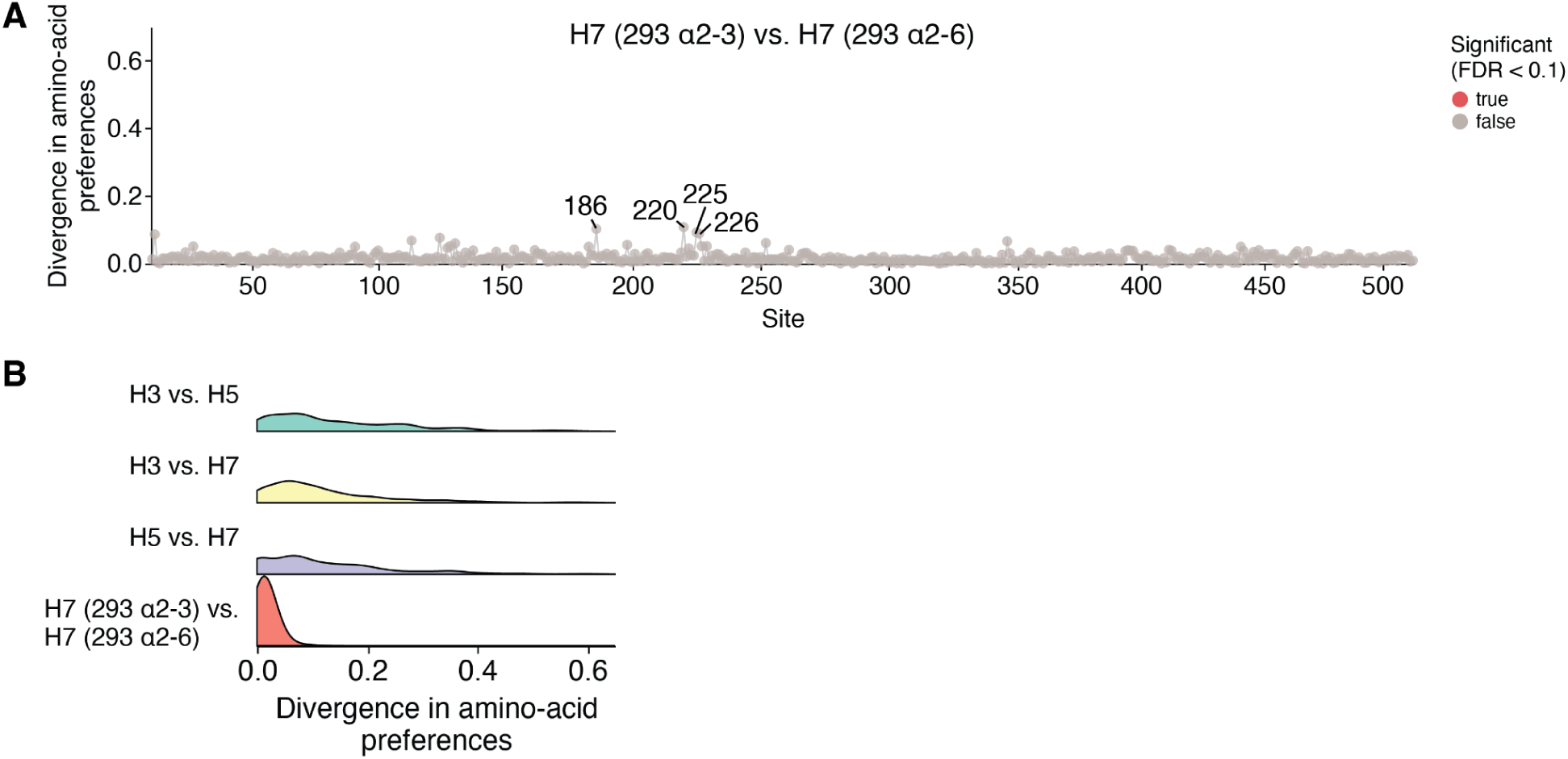
| Divergence in H7 HA amino-acid preferences for entry into 293 cells expressing α2-3 versus α2-6 linked sialic acids is minimal compared to the divergence between HA subtypes. **A)** Divergence in amino-acid preferences at each site of the H7 HA as measured in 293 cells expressing only α2-3 versus only α2-6 linked sialic acids. Note that the divergence at all sites is low compared to the cross HA-subtype comparisons in Fig. 3C, and no sites are significantly diverged (false discovery rate < 0.1). The largest differences are at sites like 186, 220, 225, and 226, which are important for receptor binding. **B)** Distributions of divergence in amino-acid preferences for all sites between subtypes compared to the distribution of divergence in H7 HA for measurement made using 293 cells expressing entry only α2-3 versus only α2-6 linked sialic acids.

**Figure S6.**
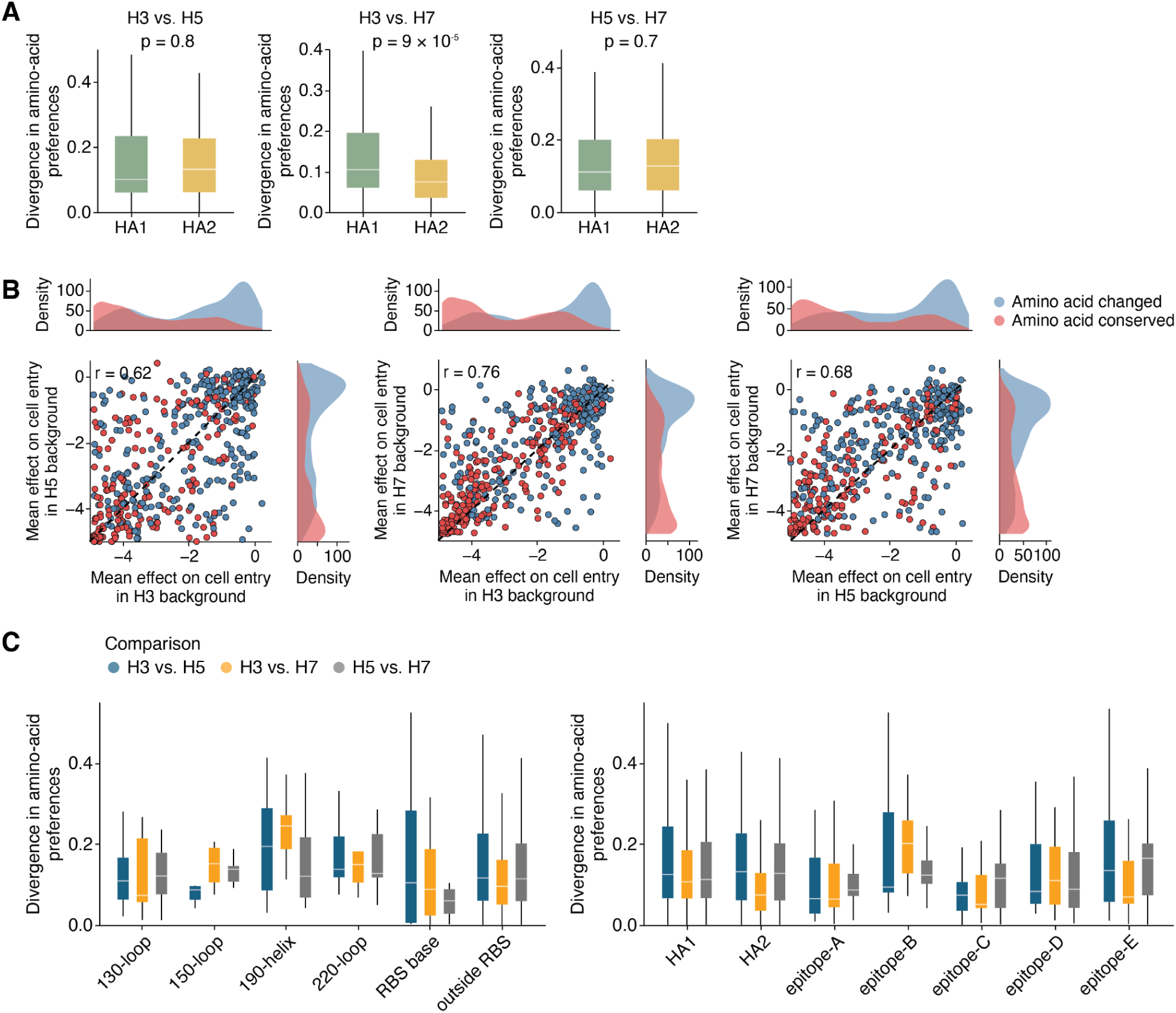
| Divergence in amino-acid preferences for different domains or regions of HA. **A)** Distributions of divergence in amino-acid preferences between HA subtypes for sites in the HA1 or HA2 domains of HA. In H3-H7, HA2 sites show significantly lower divergence compared to HA1. **B)** Correlation of the mean effect of mutations at each site between each pair of HA subtypes. Sites where the wildtype amino-acid identity differs between subtypes are colored blue, while sites with conserved wildtype amino-acid identities are red. **C)** Distributions of divergence in the amino-acid preferences by receptor binding pocket region (left) and antigenic region (right) across each HA comparison.

**Figure S7.**
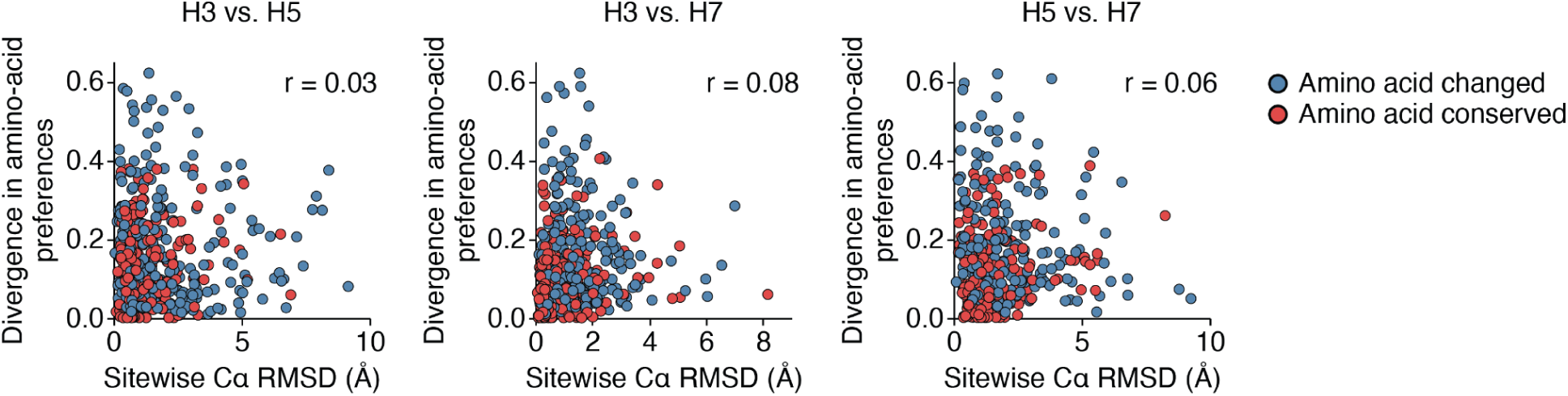
| Structural deviation is a poor predictor of divergence in amino-acid preferences at a site. Correlation between divergence in amino-acid preferences and the Cα root mean square deviation at each site in the structurally aligned backbones of HA1 and HA2 across pairwise HA comparisons. Sites where the wildtype amino acids changed between subtypes are colored blue, while conserved sites are red.

**Figure S8.**
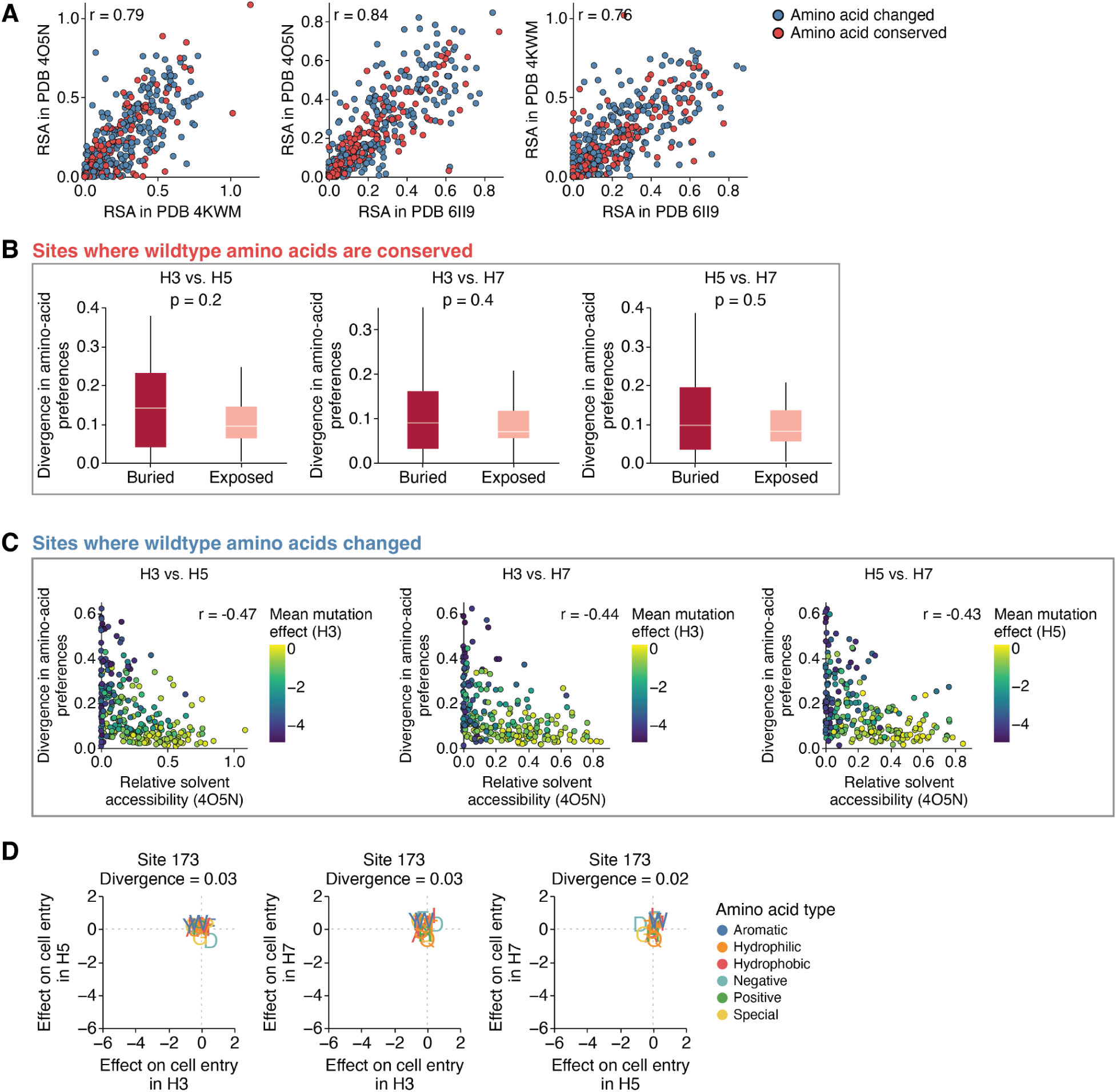
| Surface exposed sites are more mutationally tolerant and display less divergent amino acid preferences across HAs. **A)** Correlation between relative solvent accessibility values at structurally aligned sites between HAs. These values quantify how exposed or buried a site is, and the correlation is strong (*r* > 0.75) because the HA structures are conserved. PDB accessions are 4O5N (H3), 4KWM (H5), 6II9 and (H7). **B)** Among sites where the wildtype amino-acid identity is conserved (red boxplots in Fig. 4A), there is no significant difference in divergence in amino-acid preferences at buried versus exposed sites. **C)** Among sites where the wildtype amino-acid identity changed (blue boxplots in Fig. 4A), there is a negative correlation between divergence in amino-acid preferences and relative solvent accessibility. Sites are colored by mean mutation effect on entry in the indicated HA background, showing that sites that are more exposed tend to tolerate mutations better. **D)** Correlations of effects of mutations on cell entry between HAs at the exposed site 173, showing how all measured mutations are tolerated across HAs at this site. Two-sided Mann-Whitney-U test was used for all significance testing.

**Figure S9.**
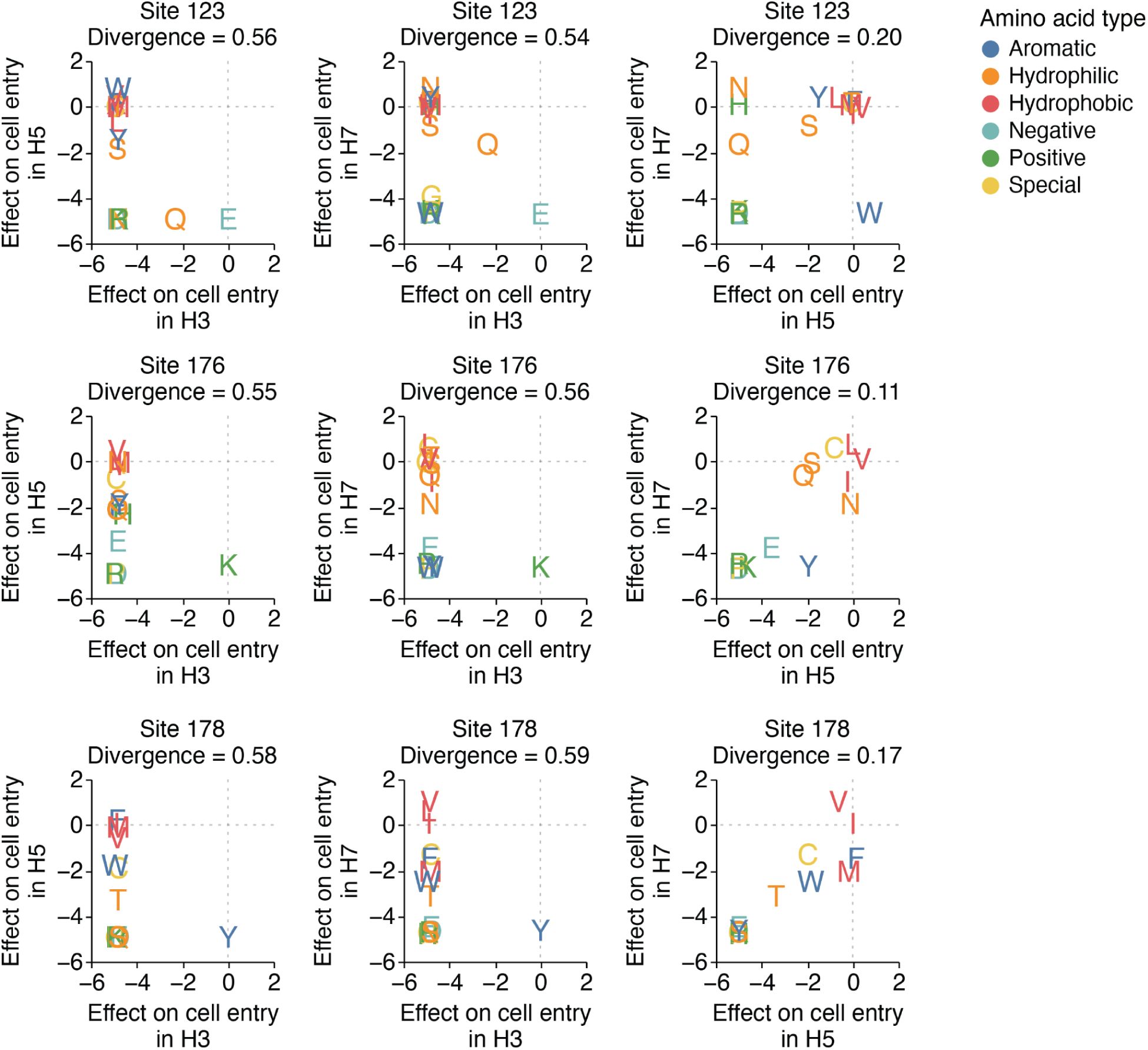
| Comparison of mutation effects across H3, H5, and H7 at buried sites 123, 176, and 178. Correlation of mutation effects on cell entry as measured by deep mutational scanning between pairs of HAs. The letters show the effect of mutation to each amino acid in each of the two indicated HA homologs. The mutation effects at these sites are relatively similar between H5 and H7, but both are highly diverged from H3 HA.

## Methods

### Code and data availability

The computer code for the deep mutational scanning of the H7 HA described in this manuscript is available on GitHub at https://github.com/dms-vep/Flu_H7_Anhui13_DMS and can be viewed interactively at https://dms-vep.org/Flu_H7_Anhui13_DMS/ via GitHub Pages. The quality filtered measurements of mutation effects on cell entry for the H7 HA are available in tabular format at https://github.com/dms-vep/Flu_H7_Anhui13_DMS/blob/main/results/summaries/entry_all_cells.csv.

A single CSV with the effects of mutations at each site on entry for the H7 HA measured in this study as well as the previously measured values for the H5 and H3 HAs^28,29^ is at https://github.com/jbloomlab/ha-preference-shifts/blob/main/results/combined_effects/combined_mutation_effects.csv. The computer code for comparing mutation effects across H3, H5, and H7 HAs is on GitHub at https://github.com/jbloomlab/ha-preference-shifts and an interactive plot showing divergence in amino-acid preferences across the three HAs is at https://jbloomlab.github.io/ha-preference-shifts/htmls/combined_interactive_lineplots.html.

### Biosafety

All experiments reported in this paper used lentiviral particles pseudotyped with HA and NA. These particles encode no viral genes other than HA and so are only able to undergo a single round of cell entry, meaning that they are not fully infectious viruses and so are not human pathogens.

### Cell lines and media

HEK293 isogenic α2-3 and α2-6 cells were stably engineered as previously described^54,55^. The α2-3 cells are 293 cells with combinatorial knockout of ST3GAL3, ST3GAL4, ST3GAL6, ST6GAL1, and ST6GAL2 genes and knock-in of the human ST3GAL4 gene. The α2-6 cells are 293 cells with knockout of ST3GAL3, ST3GAL4, ST3GAL6, ST6GAL1, and ST6GAL2 genes and knock-in of the human ST6GAL1 gene. It is important to note that both cell lines express α2-3 and α2-6-linked sialic acids on O-linked glycans.

These cells were maintained in D10 media (Dulbecco’s Modified Eagle Medium with 10% heat-inactivated fetal bovine serum, 2 mM l-glutamine, 100 U/mL penicillin, and 100 μg/mL streptomycin). To suppress rtTA activation, 293T-rtTA cells were grown in tet-free D10, which is made with tetracycline-negative fetal bovine serum (Gemini Bio, Ref. No. 100-800) instead. Additionally, a phenol free version of D10 was used for library production and rescue. Influenza growth medium was also used during library production (IGM, Opti-MEM supplemented with 0.01% heat-inactivated fetal bovine serum, 0.3% bovine serum albumin, 100 μg/mL of calcium chloride, 100 U/mL penicillin, and 100 μg/mL streptomycin).

### Design of H7 HA deep mutational scanning libraries

Deep mutational scanning libraries were designed in the background of the HA of the A/Anhui/1/2013 (H7N9) strain. The HA gene was codon optimized for use in the lentiviral backbone, see https://github.com/dms-vep/Flu_H7_Anhui13_DMS/blob/main/library_design/pH2rU3_d1GTSS_extgag_extcppt_ForInd_Anhui13_A-Anhui-1-2013_genscript2_CMV_ZsGT2APurR.gb for a map of the lentiviral backbone plasmid encoding the HA gene. The codon optimization tool used was the Gensmart codon optimization offered by Genscript. This lentiviral backbone contains several modifications relative to commonly used lentiviral vectors. We used an extended gag sequence, as has been shown to improve genome packaging efficiency^68,69^. Under the same rationale, we used an extended cppt sequence with more of the native HIV sequence following the cppt. We also mutated the transcription start site of HIV to “TTG” from “GGG” to generate TTG-initiated transcripts more likely to be packaged than other transcription start initiation site forms^70,71^.

The library was designed to have all possible amino acid mutations to each site in the HA ectodomain and a stop codon at every other position for the first 20 positions of HA. Single mutant libraries were ordered from Twist Biosciences. Quality control data for library production by Twist is at (https://github.com/dms-vep/Flu_H7_Anhui13_DMS/blob/main/library_design/Final_QC_Report_Q-367972_for_twist_libraries.xlsx)

### H7 HA deep mutational scanning plasmid library production

The H7 library ordered from Twist was designed to have mutations to all amino acids in the 506 residue ectodomain of HA. Of the 506 sites, 12 sites failed to be properly mutagenized in the Twist library production; of the sites that failed, 2 sites were in antigenic regions and within the receptor-binding pocket. To add mutations at the missing sites, forward and reverse primers with NNG/NNC codons were designed and ordered from IDT, sequences can be found at (https://github.com/dms-vep/Flu_H7_Anhui13_DMS/blob/main/library_design/Anhui13_gen2_ha_missing_mut_primers.csv and https://github.com/dms-vep/Flu_H7_Anhui13_DMS/blob/main/library_design/Anhui13_gen2_ha_missing_site_NNS_primers.csv). Forward and reverse primer pools were made by pooling the primers at equal molar ratios for a final concentration of 5µM. The linear HA template was produced by digesting the lentivirus backbone encoding the HA (plasmid map linked above) with XbaI and NotI-HF and then gel purifying the reaction. PCR mutagenesis was then performed as previously described^27,29^ with a few changes, 9 PCR cycles were used for the mutagenesis and after the joining PCR a DpnI digest was performed for 20 minutes at 37° to remove any wildtype template. This PCR mutagenesis reaction was performed in duplicate, one for library A and one for library B, resulting in two mutagenesis pools. We barcoded the library pools generated from Twist Biosciences independently in 2 separate reactions to generate library A and library B pools to serve as biological replicates, each of which were then handled independently throughout deep mutational scanning. The two mutagenized pools, A and B, were also barcoded separately, resulting in 4 separate barcoding PCR reactions. The reactions were barcoded with primers containing a random 16-nucleotide sequence downstream of the stop codon. The lentiviral backbone was digested from the same plasmid that was used to generate the linear HA fragment (https://github.com/dms-vep/Flu_H7_Anhui13_DMS/blob/main/library_design/pH2rU3_d1GTSS_extgag_extcppt_ForInd_Anhui13_A-Anhui-1-2013_genscript2_CMV_ZsGT2APurR.gb) by incubating with XbaI and NotI-HF for 2 hours at 37° followed by 65° for 20 minutes to heat inactivate XbaI. The barcoded libraries and the digested lentiviral backbone were then run on a 0.8% agarose gel at 90V for 1 hour. The bands of the appropriate size were excised using the Nucleospin Gel and PCR Clean-up Kit (Macherey-Nagel, Cat. No. 740609.5) and then purified using Ampure XP beads (Beckman Coulter, Cat. No. A63881). All products were eluted in molecular grade water.

The barcoded libraries were then cloned into the lentiviral backbone at a 1:2 insert to vector ratio in a HiFi assembly for 1 hour at 50°C. This was done using the NEBuilder HiFi DNA Assembly kit (NEB E5520S). The reactions were then purified with Ampure XP beads and eluted in molecular grade water. The purified products were then transformed into 10-beta electrocompetent cells (NEB, Cat. No. C3020K) using a BioRad MicroPulser Electroporator (Cat. No. 1652100), shocking at 2 kV for 5 milliseconds. For the Twist library pools A and B, 5 reactions were electroporated per library. For the mutagenesis library pools, 3 reactions were electroporated per library A and B. The resulting tubes of electroporated cells were placed on a 37° shaker for 1 hour to recover. After recovery, the Twist library pools were pooled per condition (5x tubes for Library A and 5x tubes for Library B) were pooled and transferred into ∼300ml of LB + Ampicillin (final concentration at 10µg/ml). The mutagenesis pools were pooled per condition (3x library A and 3x library B) and transferred into ∼150ml of LB + Ampicillin at the same final concentration as mentioned previously. We then plated 50µl of a 1:1000 dilution of the Twist library pools and mutagenized HA pools to calculate the CFU/ml. The total number of colonies from the electroporations were as follows: Twist A 7.2e7 CFU/ml, Twist B 4.8e7 CFU/ml, Mutagenesis A 3.6e6 CFU/ml, Mutagenesis B 4.2e6 CFU/ml. The high number of colonies is necessary to ensure high diversity of the library and to prevent bottlenecking of the mutants. After shaking overnight, the plasmids were then extracted using the QIAGEN HiSpeed Plasmid Maxi Kit (Cat. No. 12662). The mutagenized HA fragments were then pooled together with the library pool generated by Twist Biosciences at a molar ratio proportional to the number of mutants in each pool so that each variant was represented equally in the final library. For every 1µg of Twist pools, 0.0285µg of mutagenized HA pools were used because long-read PacBio sequencing of the plasmid libraries showed this ratio resulted in the most even distribution of mutations across the HA for the combined libraries.

### Production of cell-stored deep mutational scanning libraries

We generated cell-stored deep mutational scanning libraries where each cell is integrated with a single copy of a barcoded HA mutant to enable rescue of the genotype-phenotype linked pseudoviruses^27–29^. VSV-G pseudotyped viruses were produced by transfecting 2x10cm plates seeded with 8 million 293T cells with the lentiviral backbone containing barcode HA deep mutational scanning libraries (10µg per plate). Lentiviral Gag/pol, Tat, and Rev helper plasmids (AddGene numbers 204152, 204154, 204153; 1.25 µg per plate), a VSV-G expression plasmid (1µg per plate, AddGene number 204156), and a plasmid expressing the matched N9 NA from A/Anhui/1/2013 (1µg per plate). BioT transfection reagent (Bioland Scientific, Cat. No. B01-02) was used according to the manufacturer’s instructions. The NA gene came from the A/Anhui/1/2013 strain. The sequence for this NA can be found at https://github.com/dms-vep/Flu_H7_Anhui13_DMS/blob/main/library_design/HDM_N9_genscript_H7N9_A_Anhui_1_2013.gb.This NA was used because it provided high pseudovirus titers. It is important to note that VSV-G pseudoviruses do not require NA or virus release or entry, the produced virions will also have HA expressed from the lentiviral backbone on their surface, so NA is necessary to prevent HA from binding the virions back to the producer cell surface. This is important for the HA library, because different HA mutants will have different expression and sialic-acid binding properties. This could lead to a different tendency of different virions to be bound on the cell surface by HA if NA is not added at a high excess. 48 hours post transfection, the supernatant was filtered through a 0.45µm syringe filter (Corning, Cat. No. 431220) and then stored at -80°C. These viruses were then used to infect 293T cells to determine the titer in transcription units (TU) per mL, which was determined by measuring the percentage of zsGreen positive cells via flow cytometry.

The VSV-G pseudoviruses were then used to infect 293T-rtTA cells (using a specific clone that had shown to yield good pseudovirus titer^27–29^ at an infection rate of 0.75% so that most transduced cells receive only a single integrated proviral genome. The MOI was then confirmed by measuring the percentage of zsGreen positive cells via flow cytometry 48 hours post transduction. After measuring the positive cells, Library A contained 37,679 variants and, and library B contained 44,053 (**Fig. S2C**).. This allowed for coverage of each barcode ∼4x times. Transduced cells were then selected using 0.75µg/mL puromycin for 1 week (fresh media with puromycin was replenished every 48 hours) leading to a population of cells where each cell contains an integrated proviral genome encoding a single barcoded variant of HA. These cells were then frozen in 20 million cells per aliquot and stored in liquid nitrogen until further use.

### Rescue of HA and VSV-G expressing pseudovirus libraries

To rescue HA expressing pseudoviruses from the integrated cells, 150 million cells were plated in 5-layer flasks in phenol-free tetracycline-free D10 supplemented with 1µg/mL of doxycycline to induce HA expression from the integrated genomes. The next day, each flask was transfected with 40µg of each helper plasmids encoding Gag/Pol, Tat, and Rev (AddGene numbers 204152, 204154, 204153), 15µg of plasmid expressing human airway trypsin-like protease (https://github.com/dms-vep/Flu_H7_Anhui13_DMS/blob/main/library_design/HDM_HAT.gb) to activate the HA for membrane fusion, and 15µg of the plasmid expressing NA. BioT transfection reagent was used according to the manufacturer’s instructions. 16 hours post transfection, the pheno-free tetracycline-free media was aspirated off, and 150mL of fresh Influenza Growth Media with 1µg/mL of doxycycline was added. This low serum media is necessary due to the non-specific inhibitors in FBS that can inactivate HA and interfere with pseudovirus infection. 32 hours after the media swap, the supernatant was then filtered through 0.45μm SFCA Nalgene 500 mL Rapid-Flow filter unit (Cat. No. 09-740-44B). Filtered supernatant was then concentrated using 100K MWCO Pierce protein concentrators (Thermo Fisher Cat. No. 88537) that were spun down at 1500xg at 4° for 1 hour. After centrifugation, supernatant was discarded and the concentrated virus was pooled and aliquoted. The estimated titers were ∼3.2e6TU/mL. 1mL aliquots of the concentrated library viruses were frozen and stored at -80°C until further use.

To rescue VSV-G expressing pseudoviruses from integrated cells, 20 million cells were plated in 15cm plates in phenol-free tetracycline-free D10. The next day, each plate was transfected with 6.75µg of each helper plasmids expressing Gag/Pol, Rev, and Tat, 3µg of plasmid expressing NA, and 6.75µg of plasmid expressing VSV-G. 48 hours post transfection, the supernatant was filtered through a 0.45µM SFCA 500mL Rapid-Flow filter unit and concentrated using 100K MWCO Pierce protein concentrators. Aliquots of the concentrated VSV-G expressing pseudoviruses were frozen at -80°C for use in linking mutations to barcodes and cell entry experiments.

### Long-read PacBio sequencing for variant-barcode linkage

1e6 293T cells were plated per well in a 6-well plate coated with Poly-L-Lysine to help with cell adhesion. The next day, 15 million TU’s of VSV-G expressing pseudoviruses that were rescued from cell-stored deep mutational scanning libraries were used to infect the cells across 3 wells. 12 hours post infection, the non-integrated reverse-transcribed lentiviral genomes were recovered by minipreping the 293T cells using a QIAprep Spin Miniprep Kit as previously described^27–29^. Eluted genomes were split into 2 separate PCR reactions, where each reaction added a unique tag for detecting strand exchange, which could prevent correct mutation-barcode linkage. The resulting set of PCR reactions were then pooled together, and another round of PCR was performed. In each PCR round, we minimized the number of reaction cycles to limit strand exchange. The reactions were then purified with Ampure XP beads and eluted in elution buffer.Long read sequencing was then performed using a PacBio Sequel IIe machine. The PacBio circular consensus sequences (CCSs) were then aligned to the target amplicon (https://github.com/dms-vep/Flu_H7_Anhui13_DMS/blob/main/data/PacBio_amplicon.gb) using alignparse^72^, and consensus sequences for HA variants associated with each barcode were determined by requiring at least 2 CCSs per barcode. The final barcode-variant lookup table can be found at https://raw.githubusercontent.com/dms-vep/Flu_H7_Anhui13_DMS/refs/heads/main/results/variants/codon_variants.csv. Library A had 37,679 variants and library B had 44,053 variants (**Fig. S2C**).

### Mutation effects on cell entry in α2-3 and α2-6 linked sialic acids expressing cells

To measure effects of mutations on cell entry, 293 cells expressing primarily α2-3 or α2-6 linked sialic acids were used. Additionally, we created an equal pool of the α2-3 or α2-6 linked sialic acid expressing cells, and measured cell entry on this equal pool as well. These cells were then either infected with libraries pseudotyped with VSV-G envelope (produced as described above) or HA pseudotyped libraries. For HA pseudotyped libraries, the α2-3, α2-6, or an α2-3/α2-6 cell pool were plated in the presence of 2.5µg/ml of amphotericin B a day before (amphotericin B was shown to increase library virus titers), and infections were performed in the presence of 1µM oseltamivir to inhibit NA and prevent cleavage of receptors on target cell surface. For VSV-G pseudotyped library infections approximately 8 million transcription units were used and for the H7 pseudotyped library infections approximately 3.2 million transcription units were used. Because the initial titers on α2-3 vs. α2-6-linked sialic acid cells were similar, we were able to infect both cell lines with the same amount of virus. After infecting the cells with the viruses, the plates were spun in a 30° centrifuge for 1 hour at 300g to maximize infection rate. At 12 to 15 hours postinfection, cells were collected, nonintegrated lentiviral genomes were recovered using QIAprep Spin Miniprep Kit (Qiagen, cat. no. 27104), and amplicon libraries for barcode sequencing were prepared as previously described ^27–29^.

Cell entry effects of each mutation in the library were calculated as previously described^28^. Specifically, cell entry scores for each mutational variant were calculated as the log enrichment ratio: log_2_ ([n^v^_post_ / n^wt^_post_]/[n^v^_pre_ / n^wt^_pre_]), where n^v^_post_ is the count of variant *v* in the H7-pseudotyped infection (postselection condition), n^v^_pre_ is the count of variant *v* in the VSV-G-pseudotyped infection (preselection condition, and n^wt^_post_ and n^v^_pre_ are the counts for wildtype variants). Positive cell entry scores indicate that a variant is better at entering the cells compared to the unmutated parental HA, and negative scores indicate entry worse than the unmutated HA.

To calculate the mutational-level cell entry effects, a sigmoid global-epistasis^53^ function was fitted to variant entry scores for both the single- and multiple mutated variants after truncating the values at a lower bound of -5, using the *multi-dms* software package^52^. The rationale for fitting the global epistasis models is that some of the variants have multiple HA mutations, and so the model enables deconvolution of the mutation effects from these multiple mutated variants as well as just the single mutants. The mutation effects from the global epistasis model are nearly identical to those measured directly in just the single mutant variants, but the global epistasis model provides more coverage of mutations due to inclusion of the multi-mutant variants; for instance, see the plot produced by code cell [8] of https://dms-vep.org/Flu_H7_Anhui13_DMS/notebooks/avg_func_effects_293_2-6_entry.html. For the final reporting, we took the median of the estimated functional effect of each mutation across all the replicas and libraries. An interactive heatmap showing the effect of each mutation on cell entry on α2-3, α2-6, and an equal pool of both cells is at (https://dms-vep.org/Flu_H7_Anhui13_DMS/cell_entry.html). For the mutation effects, values of zero mean no effect on cell entry, negative values mean impaired cell entry, and positive values mean improved cell entry. For plots in the paper, we show only mutations observed in average of at least 2 barcoded variants per library and measured in both biological library replicates.

### Structural alignment of H3, H5, and H7 HAs

Experimentally determined HA protein structures were obtained from the Protein Data Bank (accession IDs 4O5N for H3, 4KWM for H5, and 6II9 for H7). Foldmason^73^ was used to structurally align the HA1 and HA2 domains of the three HAs. Separate alignment of the domains was necessary due to the relative shift in HA1/HA2 orientation shown in **Fig. S1**. The alignment results are at:

- HA1: https://github.com/jbloomlab/ha-epistasis/blob/main/results/foldmason_alignment/chain_A/result_aa.fa
- HA2: https://github.com/jbloomlab/ha-epistasis/blob/main/results/foldmason_alignment/chain_B/result_aa.fa

### Jensen-Shannon divergence in amino-acid preferences

For reasons described in the main text, amino-acid preferences are more suitable for comparing effects of mutations across HAs. Therefore, for each site 𝑟 and amino-acid identity 𝑎, we converted the mutation effect on cell entry 𝑥_𝑟,𝑎_ to an amino-acid preference by:

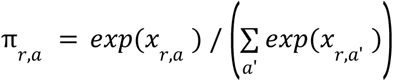

where π_𝑟,𝑎_ is the preference of site 𝑟 for amino-acid identity 𝑎, and the sum in the denominator is over all amino-acid identities 𝑎’ at that site. These amino-acid preferences can be interpreted as the probability of observing the amino-acid identity 𝑎 at a site 𝑟 in an evolutionarily equilibrated alignment evolving according to the single-mutant effects measured in the deep mutational scanning^74,75^.

To compare the amino-acid preferences between HAs, we quantified the Jensen-Shannon divergence (JSD) between their amino-acid preference probability distributions at each site. Let 𝑃 be the amino-acid preferences at a site in one HA and Q be the amino-acid preferences at the same site in another HA. Then, the JSD at the site is the average Kullback-Leibler divergence (KLD) of P and Q from their per-index-mean vector 𝑀:

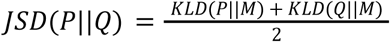

where the KLD is calculated as

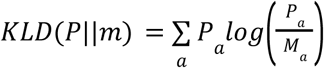

We calculated the Jensen-Shannon divergence between HAs at every site where there were at least 10 amino acids measured in both HA backgrounds.

### Significance testing for divergent amino-acid preferences between HAs

We implemented a simulation approach to test for significant divergence in amino-acid preferences between HAs. Under the null hypothesis that the true mutation effects (and therefore preferences) are identical across HAs and that any divergence is due only to measurement uncertainty, we generated a simulated null distribution for the JSD at each site.

For a given background X (e.g., H3), we drew two independent replicate measurements of each mutation effect by 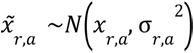, where 𝑥_𝑟,𝑎_ is the measured mutation effect and σ_𝑟,𝑎_ is the standard deviation across replicate measurements made for that HA (note that all cell entry effects for all three HAs were made in replicate with this paper typically just reporting the median value of the replicate measurements for each mutation effect). This standard deviation is calculated as a typical population standard deviation using 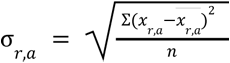, where 𝑥_𝑟,𝑎_ is the measured mutation effect in replicate 𝑟, 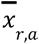 is the average mutation effect across replicates, and 𝑛 is the number of replicates. For each site, the sampled mutation effects across amino acids were converted into two amino-acid preference vectors, and the JSD between these vectors was computed. This procedure was repeated 1000 times to yield a null distribution 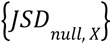. The same procedure was performed for background Y (e.g., H5) to obtain 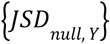. To account for potentially different levels of noise in the two backgrounds, the final null distribution was defined as the elementwise average of the two distributions, 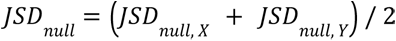.

For each site, we computed an empirical one-sided p-value as the fraction of null JSD values greater than or equal to the observed JSD. We controlled for multiple testing across sites by the Benjamini-Hochberg false discovery rate (FDR) procedure and considered sites as significant at FDR < 0.1.

Note this approach makes two assumptions about the null distribution. First, it assumes Gaussian noise in log-space. Second, it assumes the standard deviations can be faithfully derived from 2-4 replicate measurements. Therefore, a limitation is that these standard deviation estimates could be noisy, leading to false positives and negatives at sites with modest divergence values. However, clear separation between significant and non-significant sites (when divergence > 0.2) demonstrates that sites with larger divergence are robustly called significant despite potential uncertainty in the standard deviation estimates (**Fig. 3C**).

### Structural analysis

UCSF ChimeraX v.1.10.1^76^ was used for structural analysis and visualizations. In **Fig. S7**, we analyzed structural deviation between aligned HAs. The structures were aligned by Foldmason as described above and the distances between corresponding Cα atoms were computed by the rmsd function in ChimeraX. In **Fig. S8**, we analyzed the surface accessibility of sites in HA. The biological assemblies of each HA were used as input for the Define Secondary Structure of Proteins (DSSP) program^77^ to calculate the absolute solvent accessibility values of each HA site. These values were converted to relative solvent accessibility values based on the maximum allowed solvent accessibilities of residues reported in Tien et al. 2013^15^. In this work, we define a site as buried if its relative solvent accessibility < 0.2.

## Notes

https://github.com/jbloomlab/ha-preference-shifts

https://dms-vep.org/Flu_H7_Anhui13_DMS/

https://jbloomlab.github.io/ha-preference-shifts/

### Summary of Updates

Minor updates to text to clarify methods and add a new supplementary figure, as well as a bit of new discussion text.

